# Clathrin is an Intrinsic Driver of Membrane Fission

**DOI:** 10.64898/2026.03.05.709932

**Authors:** Nicoletta Bouzos, Samuel L. Foley, Ariadni Potamianos, Margaret E. Johnson, Wade F. Zeno

## Abstract

Clathrin-mediated endocytosis is a primary internalization pathway where adaptor proteins recruit clathrin and bend the plasma membrane to form clathrin coated vesicles. Since clathrin does not bind membranes directly, its contributions to membrane remodeling remain unclear. Using an in vitro system that recruits clathrin directly to lipid membranes, we show that clathrin alone can drive end-to-end membrane budding and vesicle fission. Surprisingly, stronger clathrin assembly (i.e., higher membrane association) yields weaker membrane fission, indicating that membrane remodeling depends not simply on how much clathrin is recruited, but on how the resulting lattice is assembled. Brownian dynamics simulations, together with experimentally measured shifts in clathrin cage diameter, suggest that this counterintuitive behavior arises at least in part from assembly-dependent changes in intrinsic lattice curvature. This framework extends to adaptor-mediated recruitment, with clathrin either assisting or opposing membrane fission in an adaptor-dependent manner. Together, these findings support a mechanism by which coat protein mechanics regulate membrane remodeling.

## INTRODUCTION

Internalizing cargo from the plasma membrane requires the formation and fission of membrane vesicles through essential pathways such as clathrin-mediated endocytosis (CME), a major cellular uptake pathway for nutrients, receptors, and signaling molecules in cells (*1, 2*). During CME, clathrin triskelia do not work alone – they are recruited to the membrane by adaptor proteins to assemble into lattices that stabilize clathrin-coated vesicles (*1, 3*). Clathrin triskelia are composed of three heavy chains and three light chains (*4*). While the light chains primarily regulate cage assembly (*5–7*), the heavy chains play a central role in CME: the N-terminal domain of the clathrin heavy chain interacts with adaptor proteins that are themselves associated with the membrane(*8, 9*). Through these interactions, clathrin is positioned and organized at sites of endocytosis.

The CME network consists of dozens of adaptor proteins including AP2, AP180/CALM, epsin1, and amphiphysin1(*10*). These proteins bind to phospholipids with PI(4,5)P2 headgroups and recruit clathrin through interactions with its N-terminal domain (*11–15*). Many of these adaptor proteins, including epsin1, amphiphyisn1, and AP180, have been shown to bend membranes through mechanisms such as steric crowding, amphipathic helix insertion, and BAR domain scaffolding (*13, 16, 17*). Due to simultaneous contributions of these adaptor proteins, the intrinsic contributions of clathrin in membrane remodeling have remained difficult to isolate, leaving open the question of whether clathrin itself plays a role in driving membrane fission (*18*). Recent work has provided evidence that clathrin may play a direct role in membrane fission (*19*). Experiments using nanopatterned substrates that impose nanoscale membrane curvature showed that externally induced bending can rescue vesicle formation when clathrin is depleted, suggesting that curvature progression is an essential function of clathrin (*20*). In separate studies, artificial membrane targeting of clathrin to mitochondria induced vesiculation (*21*), and high-resolution live-cell imaging revealed clathrin accumulation at membrane necks during pore constriction (*22*). Previous work has also pointed to clathrin playing a role in modulating vesicle scission(*23*). However, since clathrin does not bind membranes directly, these systems relied on clathrin that was recruited or anchored through additional factors, making it difficult to disentangle the intrinsic mechanical properties of clathrin from contributions of its membrane-associating partners. Therefore, these studies highlight the challenge of defining clathrin’s direct mechanical contribution to membrane bending and emphasize the need for experimental systems capable of isolating clathrin from its adaptor proteins.

To remove this convolution, we used a synthetic reconstitution system in which histidine-tagged clathrin binds directly to Ni^2+^ containing lipids via its N-terminal domain (Fig. 1a,b). This recruitment strategy orients clathrin on the membrane in its physiological conformation (*4*) and bypasses the need for adaptor proteins, allowing us to examine clathrin-membrane interactions, lattice assembly, and curvature generation under controlled biochemical conditions. The recombinant clathrin used here assembles indistinguishably from clathrin purified from bovine brain (*24*), indicating that the histidine tag does not alter assembly behavior.

**Figure 1:**
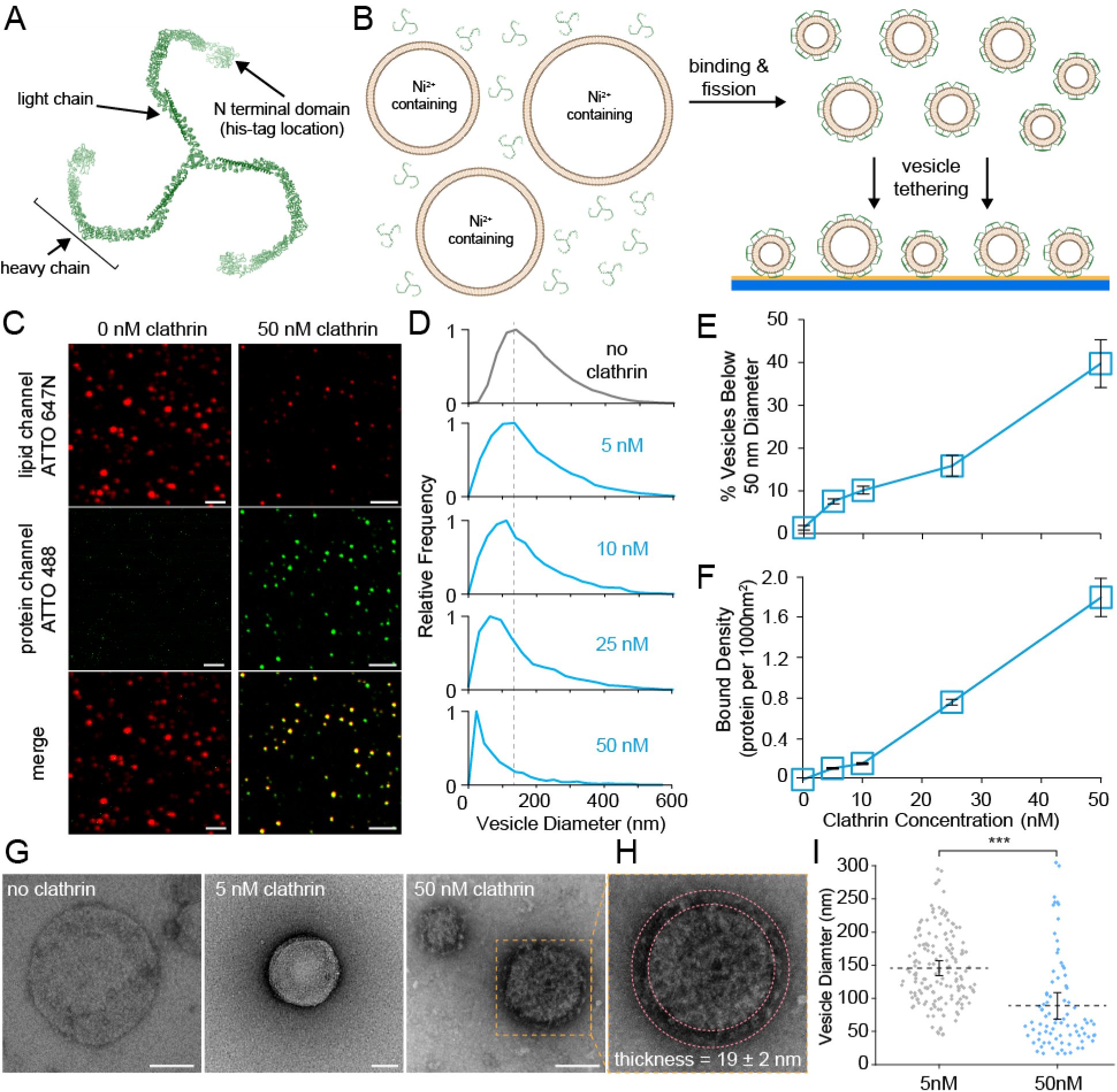
Clathrin induces membrane fission in the absence of adaptor proteins. **(A)** Structure of individual clathrin triskelion (PDB: 3IYV). **(B)** Schematic of histidine-tagged clathrin incubated with Ni^2+^ containing vesicles, leading to binding and fission. Fission products with bound clathrin were tethered to a glass coverslip (blue) that has been functionalized with biotinylated-PEG and NeutrAvidin (yellow) and analyzed for protein size and binding via fluorescence microscopy. **(C)** Representative fluorescent images of tethered vesicles and bound clathrin. Vesicle composition is 88.5 mol% DOPC, 10 mol% DGS-NTA-Ni^2+^, 1 mol% DPEG-15-biotin, and 0.5 mol% DPPE-ATTO-647N and clathrin was labeled with ATTO-488. Scale bars = 2 µm. **(D)** Histograms of vesicle diameters from confocal microscopy. No clathrin: n = 21,891 vesicles; 5 nM: n = 9,786 vesicles; 10 nM: n = 8,202 vesicles; 25 nM: n = 4,344 vesicles; 50 nM: n = 2,215 vesicles. Dashed line represents the midline of the size distribution for the control condition. **(E)** Summary of fission products from tethered vesicle experiments, expressed as the percent of vesicles in the population with diameters below 50 nm. Error bars = mean ± SD; n = 15 imaging frames. **(F)** The average number of membrane-bound clathrin triskelia per 1,000 nm^2^ of membrane surface area as a function of clathrin concentration. Error bars = 99% CI. For **(C-F)** similar results were obtained from n = 5 biological replicates. **(G)** Negative stain TEM micrographs of 200 nm extruded vesicles (90 mol% DOPC, 10 mol% DGS-NTA-Ni^2+^) before exposure to clathrin and after incubation with 5 or 50 nM clathrin. Dashed box indicates zoomed region to the right. Scale bars = 100 nm. **(H)** Zoomed in image of vesicle with assembled clathrin coat on the outer layer. Dashed concentric circles denote the boundaries of the assembled clathrin lattice used for determining coat thickness. The error represents the SD of the mean. Scale bar = 100 nm. Similar micrographs were obtained across n = 3 biological replicates. **(I)** Measured diameters of vesicles incubated with either 5 nM or 50 nM of clathrin prior to TEM imaging. Dots indicate individual vesicle measurements, and dashed lines represent the mean of the population. Error bars = 99% CI. P value - two-tailed, un-paired student’s t test. * = p <.05, ** = p <.01, *** = p <.001.

Using quantitative fluorescence microscopy and electron microscopy, we tested whether clathrin alone can induce membrane fission and how this behavior depends on lattice assembly. We modulated clathrin assembly by varying bulk solution pH and the inclusion of the divalent cation Ca^2+^, thereby altering the extent of polymerization and, consequently, the mechanical behavior of the coat. We further examined how clathrin-mediated membrane remodeling is tuned through adaptor proteins. To our surprise, enhancing clathrin assembly and membrane association reduced membrane fission efficiency, an effect that was propagated through adaptor proteins, with clathrin either opposing or enhancing adaptor driven fission. Together, these findings reveal that clathrin lattices are highly tunable in driving remodeling of biological membranes, capable of switching into highly curved, fission-inducing spherical coats dependent on adaptors and chemical environment. These results provide a general framework for proteins that coat and remodel biological membranes.

## RESULTS

### Clathrin drives membrane fission in the absence of adaptor proteins

To determine whether clathrin alone can directly remodel vesicles in the absence of adaptor proteins, we measured changes to vesicle size distributions after incubating histidine-tagged clathrin with Ni^2+^ containing vesicles (Fig. 1a,b). Vesicles were prepared by extrusion and tethered to glass coverslips for fluorescence-based measurements using a previously developed tethered-vesicle fluorescence microscopy assay (*25, 26*) (Fig. 1b). Vesicle diameters were calibrated by correlating lipid fluorescence intensity with dynamic light scattering measurements, enabling quantitative conversion of fluorescence signals into vesicle size distributions (Fig. S1). Fluorescent labeling of clathrin with ATTO-488 allowed simultaneous measurement of membrane binding (Fig. S2) and vesicle remodeling at the single-vesicle level. Incubation of vesicles (10 μM total lipid concentration) with 50 nM clathrin led to a marked reduction in lipid fluorescence intensity, indicating a decrease in vesicle diameter (Fig. 1c). Additionally, clathrin fluorescence was strongly colocalized with vesicles, confirming membrane association. Sequentially increasing the bulk clathrin concentration from 0-50 nM progressively shifted the vesicle diameter distribution toward smaller sizes, with the peak decreasing from approximately 160 nm in the absence of clathrin to approximately 50 nm at 50 nM clathrin (Fig. 1d). Consistent with this shift, the fraction of vesicles with diameters below 50 nm increased monotonically from nearly 0% to approximately 40% (Fig. 1e). Notably, this emergence of sub-50 nm diameter vesicle populations was positively correlated with an increase in clathrin binding, which reached an average value of approximately 1.8 triskelia per 1000 nm^2^ of membrane surface area (Fig. 1f, S3), consistent with reported triskelia densities on clathrin coated vesicles (*27, 28*).

Transmission electron microscopy (TEM) provided independent confirmation of these observations (Fig. 1g). At an elevated clathrin concentration of 50 nM, lattice formation could be visualized on the surface of vesicles while no such assembly was observed at 5 nM clathrin concentration. The thickness of these lattices varied between 17-21 nm (Fig. 1h, S4), matching the reported dimensions of assembled clathrin cages (*28, 29*). Increasing the clathrin concentration from 5 to 50 nM resulted in a significant reduction in TEM-measured diameters (Fig. 1i), consistent with the observations in Figures 1d-e. We note that it was not feasible to reliably quantify size distributions in the absence of clathrin as the vesicles in these protein-free samples were highly susceptible to rupture during sample preparation (Fig. S5). Together, this data demonstrates that clathrin can independently drive membrane fission.

### Clathrin assembly regulates the efficiency of membrane fission

To examine how clathrin assembly influences membrane fission, we modulated assembly by varying buffer pH (*27, 30*). Clathrin contains amphiprotic histidine residues at triskelion-triskelion binding interfaces that dictate their assembly (*31*), such that at pH 6.2 assembly is enhanced and at pH 8.3 this assembly is reduced. Fluorescent binding assays showed that decreasing pH monotonically increased clathrin recruitment to vesicles (Fig. 2a-b, S6) and enhanced binding cooperativity (Fig. 2b), consistent with strengthened protein-protein interactions and enhanced clathrin assembly. For clarity, the binding assays presented in Figures 2a-b used vesicles that were tethered before protein incubation, whereas the fission assays presented elsewhere in this work used vesicles that were tethered after protein incubation. To confirm that the observed pH dependence of binding arose from clathrin assembly, we conducted control experiments using a variant of the ENTH domain from epsin1, his-ENTHΔH0, where the H0 amphipathic helix was removed and a histidine-tag was added in its place. Across a his-ENTHΔH0 concentration range of 1 µM to 30 µM, binding exhibited no clear or strong trends with pH (Figs. S7-S8), indicating that pH-dependent binding was specific to clathrin.

**Figure 2:**
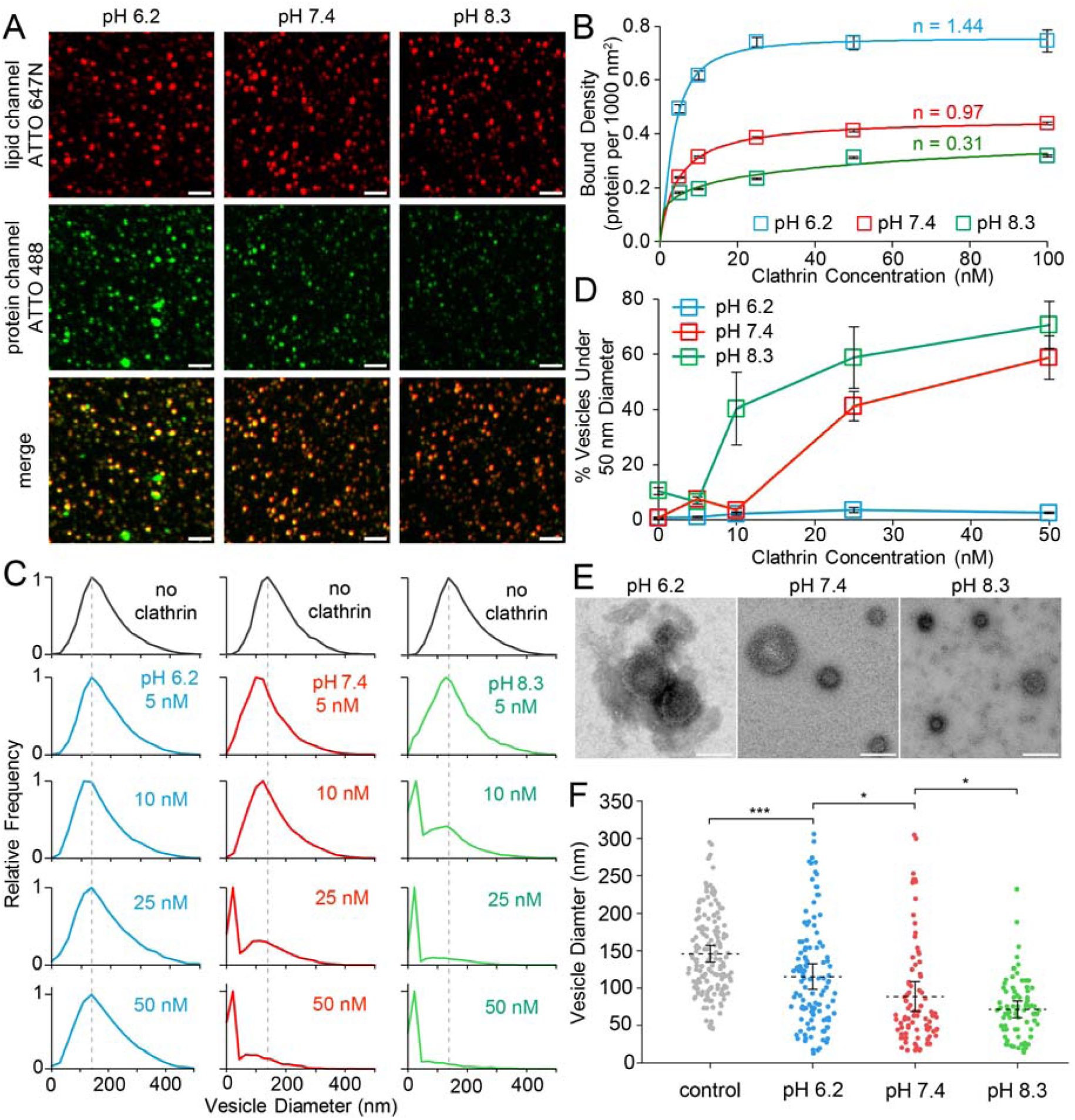
pH-dependent assembly regulates membrane fission. **(A)** Representative fluorescent confocal images of vesicles that were tethered then incubated with 50 nM clathrin at pH 6.2, 7.4, 8.3. Scale bar = 5 μm. **(B)** Number of membrane-bound proteins per 1,000 nm^2^ of membrane surface area versus concentration of clathrin for pH 6.2, 7.4, and 8.3. Raw data is composed of > 8,000 protein bound vesicles per condition. Binding isotherms for his-clathrin are fitted with Eq. 1, with cooperativity (n) values included. Error bars represent a 99% CI. Clathrin was fluorescently labeled with ATTO-488. For both **(A,B)** similar results were obtained across n = 4 independent replicates. **(C)** Histograms of vesicle diameters determined via confocal microscopy at pH 6.2, 7.4, and 8.3. n > 2,500 vesicles for each condition. For these fission experiments, vesicles and clathrin were incubated at 37°C prior to tethering and imaging. Dashed lines represent the midline of the control. **(D)** Summary of fission products from tethered vesicle experiments, expressed as the percent of vesicle diameters within the high fission product group of 50 nm or smaller. Error bars = mean ± first SD; n = 15 frames. For both **(C,D)** similar results were obtained across n = 5 independent replicates. Vesicles for **(A)** – **(D)** were composed of 88.5% DOPC, 10% DGS-NTA-NI^2+^, 1% DPEG-15-biotin, and 0.5% DPPE-ATTO-647 and were extruded through 200 nm pores. **(E)** Negative stain TEM micrographs of 200 nm extruded vesicles after incubation with 50 nM clathrin at pH 6.2, 7.4, and 8.3. Vesicles were composed of 90 mol% DOPC and 10 mol% DGS-NTA-Ni^2+^ and extruded through 200 nm pores. Scale bars = 100 nm. Similar micrographs were obtained across n = 3 independent replicates. **(F)** Vesicle diameters after incubation with 50 nM of clathrin determined from electron micrographs. Clathrin cage diameters for each condition can be found in Figure S5. “Control” corresponds to vesicles incubated with 5 nM clathrin at pH 7.4. Dots indicate vesicles, and dashed lines represent the mean of the population. Error bars = 99% CI. P value - two-tailed, un-paired Student’s t test. * = p <.05, ** = p <.01, *** = p <.001.

Clathrin drove the most potent vesicle fission at pH 8.3, whereas no appreciable membrane fission was observed at pH 6.2 across the entire protein concentration range tested (Figs. 2c-d). In control experiments where his-ENTHΔH0 was used to drive membrane fission via protein crowding, pH modulation resulted in no monotonic fission trends (Figs. S7-S8), demonstrating that pH modulated fission was unique to clathrin in our experimental setup. The inability of clathrin to drive substantial membrane fission at pH 6.2 was particularly striking, given that this condition yielded the highest surface concentration of clathrin. This result demonstrates that the architecture of clathrin assembly, rather than surface density, dictates the efficiency of membrane fission. These results were corroborated by TEM measurements, where lattice assembly was observed at all pH values (Fig. 2e) and the vesicle diameters shifted to smaller populations as pH was increased (Fig. 2f).

In addition to pH, clathrin assembly can be modulated by divalent cations such as Ca^2+^, which stabilizes clathrin-clathrin interactions by binding to the light chain and reducing electrostatic repulsion (*32*). The addition of 1 µM CaCl_2_ to our experimental buffers increased clathrin binding and cooperativity to Ni^2+^ containing vesicles across the entire clathrin concentration range tested (Figs. 3a-b, S9), consistent with enhanced assembly. Similar to the results observed with the experiments in which pH was modulated, the presence of calcium markedly reduced membrane fission, particularly at higher clathrin concentrations where membrane binding was highest (Fig. 3c,d). These results were supported by TEM data, which confirmed that clathrin coats formed under calcium addition (Fig. 3e), but vesicles incubated with calcium exhibited only minor reductions in size relative to controls (Fig. 3f). Together, these results show that calcium enhanced clathrin assembly increases membrane binding while suppressing fission, mirroring the effects observed under acidic pH conditions.

**Figure 3:**
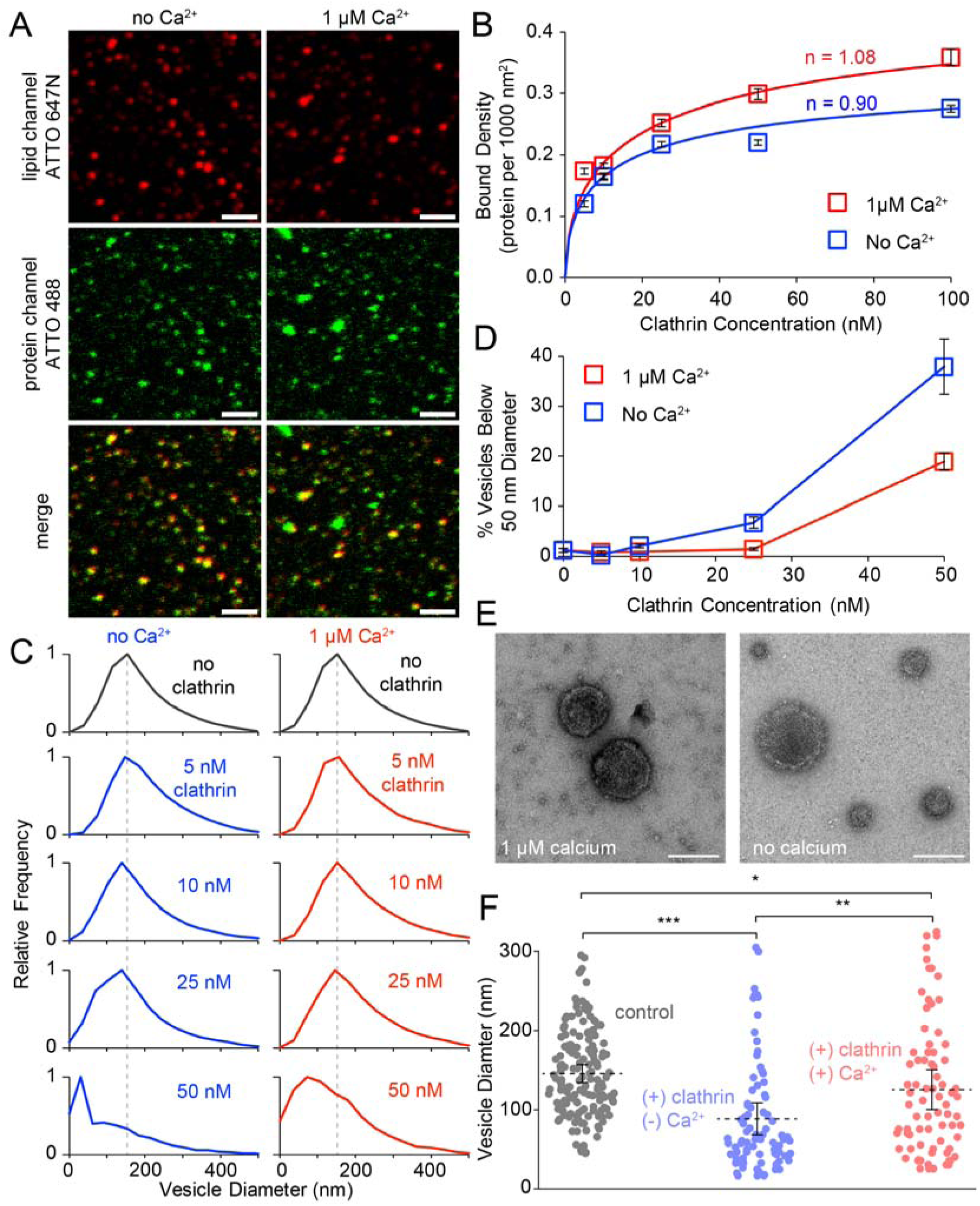
Divalent cation dependent clathrin assembly regulates membrane fission. **(A)** Representative fluorescent confocal images of vesicles that were tethered, then incubated with 50 nM clathrin in the absence or presence of 1 μM Ca^2+^. Scale bar = 2 μm. **(B)** Number of membrane-bound proteins per 1,000 nm^2^ of membrane surface area versus concentration of clathrin for vesicles incubated in the presence or absence of 1 μM Ca^2+^. Raw data is composed of > 6,000 protein bound vesicles per condition. Binding isotherms for his-clathrin are fitted with Eq. 1, with cooperativity (n) values included. Error bars represent a 99% CI. Clathrin was fluorescently labeled with ATTO-488. For both **(A,B)** similar results were obtained across n = 3 independent replicates. **(C)** Histograms of vesicle diameters determined via confocal microscopy in the absence or presence of 1 μM Ca^2+^. n > 4,500 vesicles for each condition. For these fission experiments, vesicles and clathrin were incubated at 37°C prior to tethering and imaging. Dashed lines represent the midline of the control. **(D)** Summary of fission products from tethered vesicle experiments, expressed as the percent of vesicle diameters within the high fission product group of 50 nm or smaller. Error bars = mean ± first SD; n = 10 frames. For both **(C,D)** similar results were obtained across n = 4 independent replicates. Vesicles for **(A)** – **(D)** were composed of 88.5% DOPC, 10% DGS-NTA-NI^2+^, 1% DPEG-15-biotin, and 0.5% DPPE-ATTO-647 and were extruded through 200 nm pores. **(E)** Negative stain TEM micrographs of vesicles after incubation with 50 nM clathrin. Vesicles were composed of 90 mol% DOPC and 10 mol% DGS-NTA-Ni^2+^ and extruded through 200 nm pores. Scale bars = 100 nm. Similar micrographs were obtained across n = 3 independent replicates. **(F)** Vesicle diameters after incubation with 50 nM of clathrin, determined from electron micrographs. Clathrin cage diameters for each condition can be found in Figure S5. “Control” corresponds to vesicles incubated with 5 nM clathrin at pH 7.4. Dots indicate vesicles, and dashed lines represent the mean of the population. Error bars = 99% CI. P value - two-tailed, un-paired Student’s t test. * = p <.05, ** = p <.01, *** = p <.001.

### Simulations Identify Structural and Mechanical Determinants of Clathrin-Induced Membrane Curvature

To probe how assembly-dependent changes in clathrin structure and mechanics influence curvature generation, we employed continuum membrane simulations in which elastic clathrin lattices were coupled to the membrane surface (*33, 34*) (Fig. 4a and Methods). The primary goal of this model was to test whether curvature generation is dictated by (i) the rate at which clathrin binds and assembles on the membrane, (ii) the intrinsic structural properties of triskelia as determined by the pucker angle, or (iii) the mechanical properties of assembled lattices. A key assumption of this model is that clathrin is recruited to the membrane and assembles into stable lattice structures, as observed in the experiments above. Accordingly, the model does not address the conditions that drive clathrin recruitment and stable assembly (*33*). Clathrin dissociation from assembling lattices is also not permitted, consistent with our FRAP analysis, which showed no measurable dynamic exchange over a 250-second interval (Fig. S10). We address the following question: given that clathrin lattices assemble and are capable of membrane scission under the experimental conditions tested (Figs. 1-3), how does each of these three listed properties impact curvature generation?

**Figure 4:**
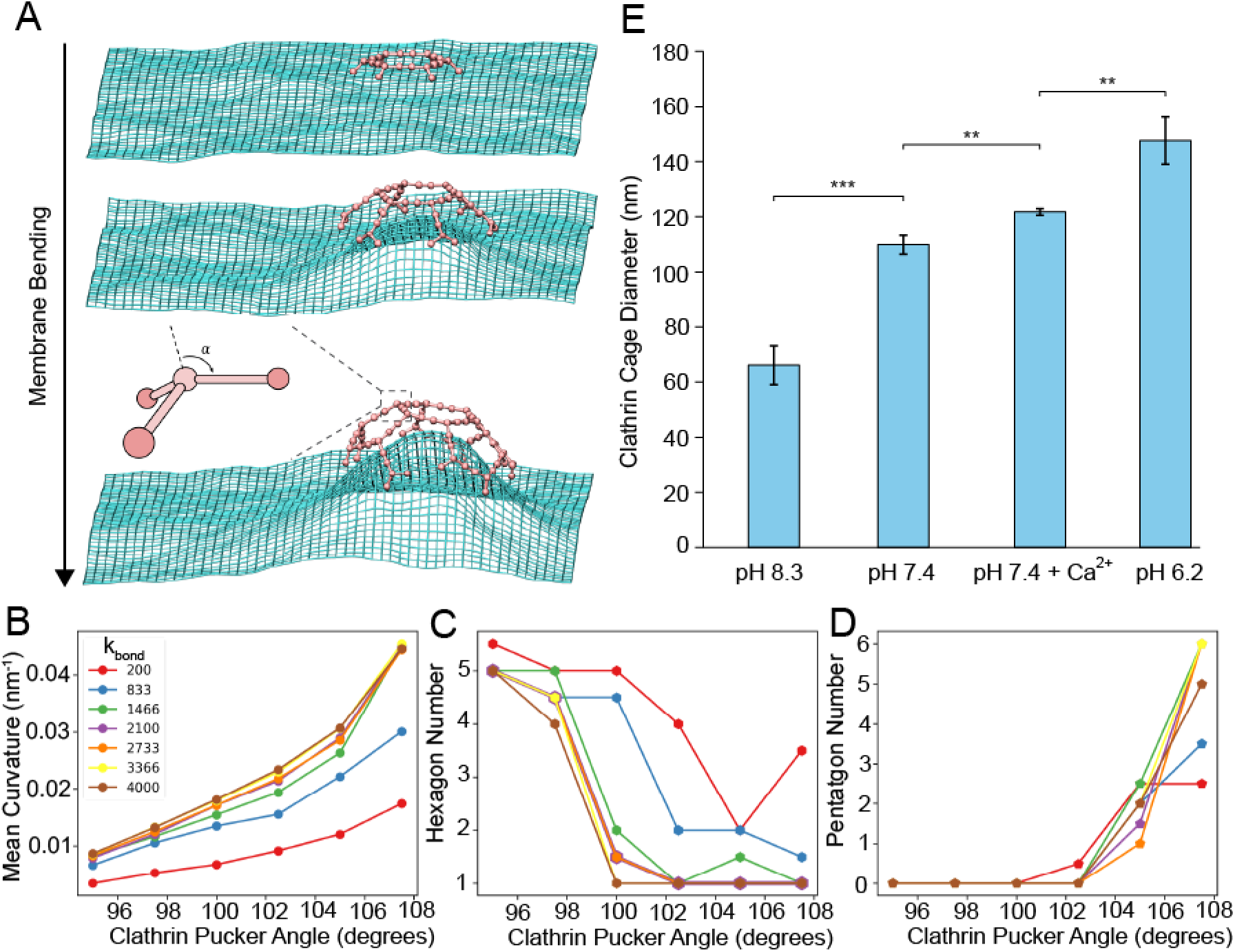
Structural and mechanical properties of clathrin govern its capacity to generate membrane curvature. **(A)** Bilayer continuum membrane model used to simulate membrane bending beneath elastic clathrin lattices. We change clathrin triskelia structure by increasing clathrin pucker angle (*a*, diagram shown in inset) and increase lattice stiffness by increasing the elastic force constant k_bond_ (units k_B_T/nm^2^ or k_B_T/rad^2^; see Methods), impacting **(B)** equilibrium lattice curvature, **(C)** number of hexagons in lattice, and **(D)** number of pentagons in lattice. **(E)** Mean hydrodynamic diameter of clathrin cages obtained from dynamic light scattering. Error bars = mean ± first SD; n = 3 measurements. P value - two-tailed, un-paired Student’s t test. * = p <.05, ** = p <.01, *** = p <.001.

In this model, individual clathrin triskelia are treated as rigid bodies propagated using standard Brownian dynamics, with each triskelion containing three binding sites that allow it to associate with neighboring triskelia and form a harmonically bonded lattice. Each triskelion is coupled to the membrane via a harmonic spring connected to its center of mass (Fig. S11). The membrane itself is parametrized by a height function, *h(x,y)* and simulated using the Fourier-space Brownian dynamics (FSBD) method of Lin and Brown (*35*), which propagates the small-gradient Helfrich functional in the over-damped regime. Because the height function representation cannot accommodate membrane scission or near-scission geometries, we focus on quantifying the membrane curvature induced during nucleation and growth; stages of clathrin coat assembly that must precede scission. The membrane bending rigidity was fixed at 20 k_B_T (*36*).

The key parameters varied across our models are the triskelion pucker angle α (Fig. 4a), which controls the curvature preference of individual triskelia and the rigidity of the assembled lattice, which is controlled by the elastic force constants *k*_bond_ (see Methods). These parameters were chosen because they are known or reasonably expected to change under different assembly conditions (*5, 6, 37–39*). Factors such as pH, divalent cations, and solvent composition modulate clathrin–clathrin interactions and may thereby alter triskelion pucker and the stiffness of the assembled lattice. The simulations assume tensionless membranes. Although the large-scale clathrin-induced deformations that lead to scission may generate membrane tension, this does not alter the qualitative conclusions drawn from our model because, to lowest order, an increase in tension appears as an effective increase in the membrane rigidity (*34*). The precise tension state of the vesicle membranes in the experiments will depend on the specific clathrin geometries and whether the strong, near-scission deformations permit water to exchange between the interior and exterior of the vesicles. Such exchange would reduce membrane tension by allowing the vesicle volume to decrease.

We first examined whether the kinetics of clathrin binding would alter membrane deformation by trapping the lattice into flatter and thus less flexible structures, as we do not allow disassembly. If the triskelia assemble more rapidly than the membrane bends in response, the lattice will tend to grow hexagonally, which may impede spherical bending. We found that altering binding kinetics over an order of magnitude did not significantly affect curvature generation, as the speed of membrane deformation exceeds that of the assembly process (Fig. S12). In contrast, varying structural parameters strongly influenced membrane deformation. Increasing the clathrin pucker angle from 96° to 108° substantially increased curvature generation for all values of lattice stiffness tested (Fig. 4b). Increasing lattice stiffness also produced a monotonic increase in curvature at fixed pucker angles. These changes correlated with increases in pentagon-to-hexagon ratios within the lattice, a geometric feature known to promote strongly curved rather than flat membrane architectures (*40*) (Fig. 4c,d). Although these simulations do not establish that more highly curved and stiffer lattices promote stronger membrane scission, they demonstrate that such lattices generate greater membrane curvature, which is a necessary prerequisite for scission of clathrin-coated vesicles. More highly curved vesicles also require fewer clathrin triskelia to complete the lattice, which could help offset the weaker assembly strength observed at pH 8.3.

In the absence of membranes, clathrin can form closed, spherical lattices in solution (*30, 41*), reflecting an intrinsic curvature preference. Therefore, to connect the simulated pucker-angle dependence to experimental outcomes, we monitored clathrin cage assembly in solution using dynamic light scattering (DLS) and quantified the resulting diameter distributions (Fig. 4e, S13). Across solution conditions known to modulate clathrin–clathrin interactions (e.g., pH and Ca²), we observed systematic shifts in cage diameter spanning approximately 50–200 nm. Qualitatively, conditions that produced smaller cages are consistent with larger triskelion pucker angles in our simulations, as increased pucker corresponds to greater intrinsic curvature and therefore smaller equilibrium cage diameters (*42*). In other words, more highly curved triskelia assemble into more tightly curved lattices, potentially increasing the bending driving force and as a result, yield smaller cages.

Importantly, the experimentally measured diameter range aligns well with the geometric predictions from our simulation framework. Simulations spanning pucker angles from 95° to 107.5° generate cage diameters of roughly 50–180 nm (*42*), closely matching the DLS distributions in Figure 4e. This agreement does not establish the pucker angle of individual triskelia, but demonstrates that the experimentally observed changes in intrinsic cage curvature fall within the range produced by varying pucker angle in the model. Moreover, the range of curvatures obtained in our simulations in Figure 4b (0.01–0.04 nm ¹) correspond to radii of curvature as high as 25 nm, consistent with our operational cutoff used in experiments to define vesicle fission products, 50 nm diameter. Taken together, these comparisons support an assembly-dependent change in intrinsic lattice curvature, consistent with a contribution from triskelion pucker, as a mechanistic link between the experimental perturbations and membrane fission.

### Clathrin inhibits amphiphysin1-mediated membrane fission

To extend our understanding beyond direct clathrin remodeling of vesicles, we next examined the more physiologically relevant question of how clathrin assembly influences membrane fission when recruited by two of its adaptor proteins, amphiphysin1 and epsin1. Amphiphysin1 has a 242-residue N-BAR domain that is known to assemble on curved membranes and drive membrane tubulation, while the full-length protein has been shown to promote membrane fission due to steric crowding from its large, 380-residue intrinsically disordered domain (*43, 44*). Amphiphysin1 contains two clathrin interacting motifs within its intrinsically disordered region that bind to the N-terminal domain of clathrin heavy chain (*45*) (Fig 5a).

**Figure 5:**
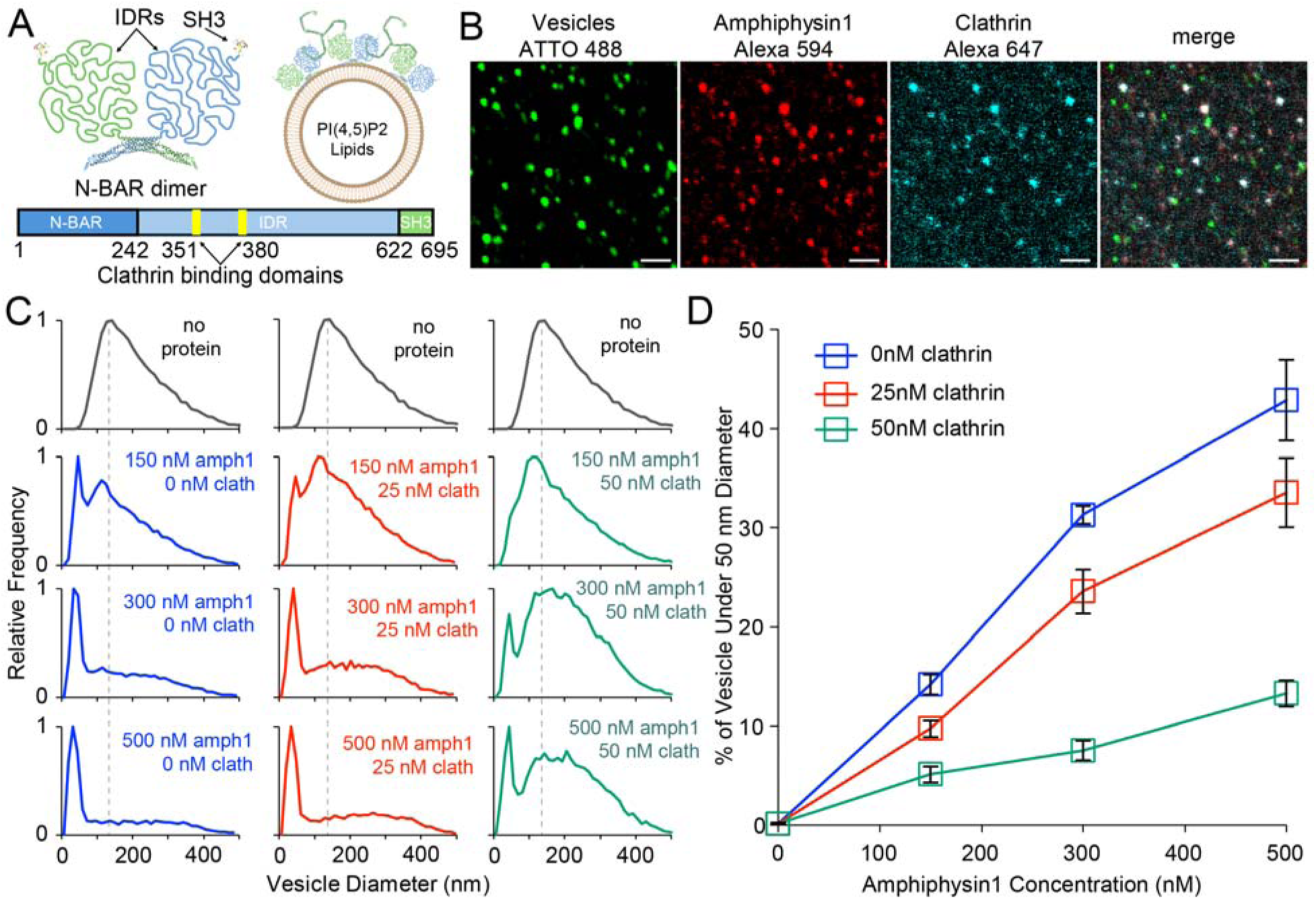
Clathrin inhibits amphiphysin1-mediated membrane fission. **(A)** Schematic of amphiphysin1’s structure, which includes the N-BAR domain (PDB: 4ATM), intrinsically disordered region, and SH3 domain (PDB: 1BB9). A schematic of clathrin recruitment to PI(4,5)P2 vesicles via amphiphysin1 is included. **(B)** Representative fluorescent confocal images of vesicles that were tethered, then incubated simultaneously with amphiphysin1 and clathrin. Vesicles were fluorescently labeled with ATTO 488-DPPE. Amphiphysin1 was fluorescently labeled with Alexa Fluor 594. Clathrin was fluorescently labeled with Alexa Fluor 647. Protein concentrations used in these images were 500 nM for amphiphysin1 and 50 nM for clathrin. Vesicles used were composed of 78.5 mol% DOPC, 15 mol% DOPS, 5 mol% PI(4,5)P2, 1 mol% DPEG-15-biotin, and 0.5 mol% DPPE-ATTO488 and extruded through 200 nm pores. Scale bars = 2 μm. Similar results were obtained across n = 3 independent replicates. **(C)** Histograms of vesicle diameters determined via confocal microscopy. n > 4,800 vesicles for each condition. For these fission experiments, vesicles, clathrin, and amphiphysin1 were incubated at 37°C prior to tethering and imaging. Dashed lines represent the midline of the distribution for the control condition. **(D)** Summary of fission products from tethered vesicle experiments, expressed as the percent of vesicle diameters within the high fission product group of 50 nm or smaller. Error bars = mean ± first SD; n = 10 frames. For both **(C,D)** similar results were obtained across n = 4 independent replicates. Vesicles for **(C)** and **(D)** were composed of 78.5 mol% DOPC, 15 mol% DOPS, 5 mol% PI(4,5)P2, 1 mol% DPEG-15-biotin, and 0.5 mol% DPPE-ATTO647N and extruded through 200 nm pores.

Three color confocal microscopy confirmed that amphiphysin1 recruited clathrin to vesicles containing 5 mol% PI-(4,5)P2 (Fig. 5b). Next, membrane fission was monitored by incubating vesicles with increasing concentrations of amphiphysin1, either in the absence or presence of clathrin (Figures 5c,d). Amphiphysin1 alone produced substantial membrane fission products as its concentration was increased to 500 nM, agreeing with previously reported results (*44*). However, the inclusion of clathrin suppressed amphiphysin1 driven fission. Increasing clathrin concentration progressively shifted vesicle populations towards larger sizes (Fig. 5c) and reduced the fraction of fission products across the amphiphysin1 concentration range tested (Fig. 5d). Furthermore, binding measurements show that clathrin enhanced amphiphysin1 association to the membrane (Fig. 5b, Fig. S14), suggesting that clathrin assembly stabilizes amphiphysin1 on the membrane in a configuration that diminishes the overall capacity of the two-protein system to drive fission.

### Clathrin enhances epsin1-mediated membrane fission

We next examined whether clathrin could regulate membrane fission of epsin1, an early-stage CME adaptor protein (*46*). Similarly to amphiphysin1, epsin1 binds to the terminal domain of clathrin via two motifs located within its C-terminal intrinsically disordered domain (*45, 47*) and has an affinity for PI(4,5)P2 lipids through its Epsin N-terminal Homology (ENTH) domain (*13*) (Fig 6a). Epsin1 has previously been shown to drive membrane fission through steric crowding of its large intrinsically disordered domain (*48*) as well as membrane insertion of the amphipathic helix within its ENTH domain (*49*). Three-color fluorescence imaging revealed that, like amphiphysin1, epsin1 bound to membranes and clathrin simultaneously (Fig. 6b).

**Figure 6:**
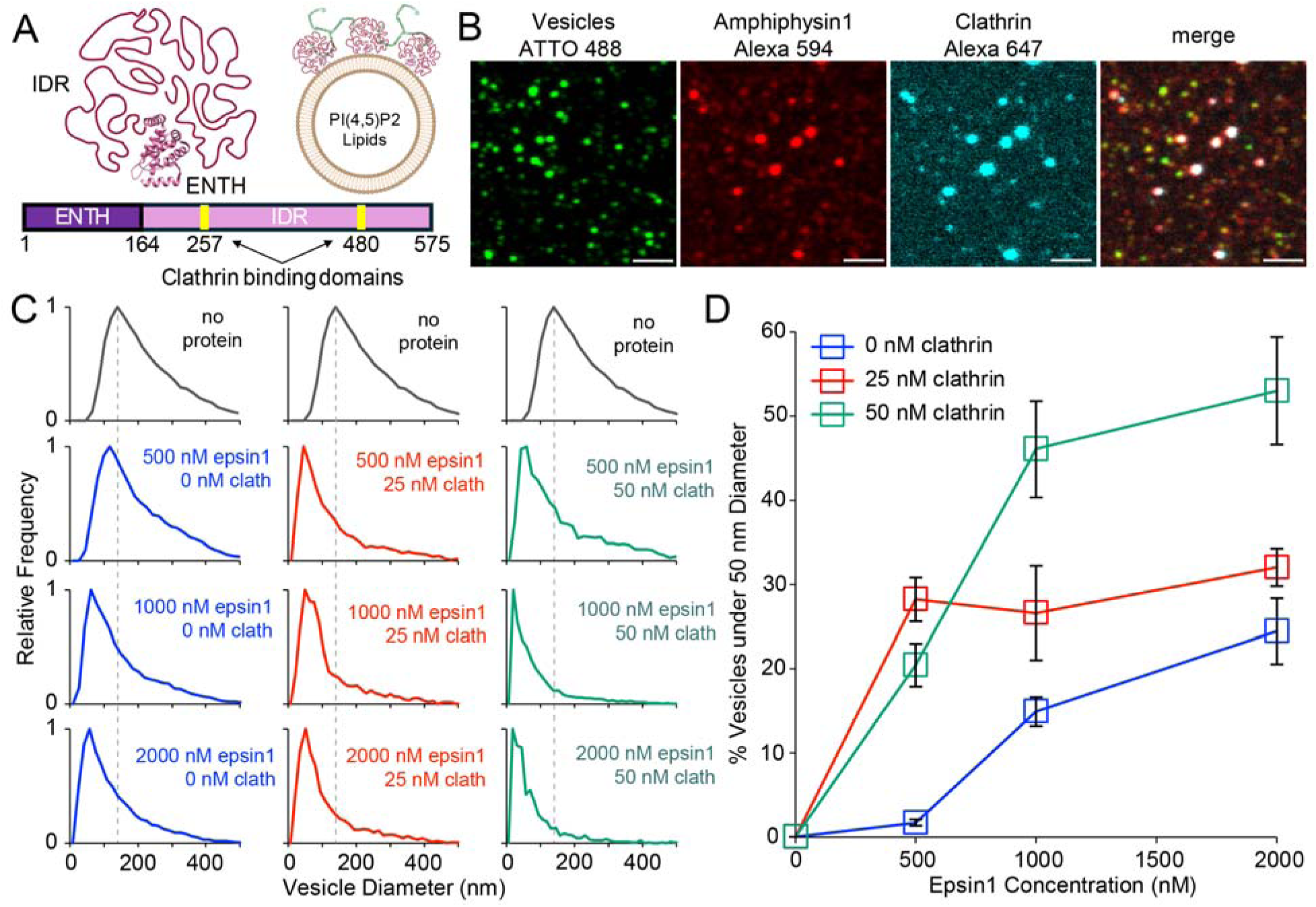
Clathrin enhances epsin1-mediated membrane fission. **(A)** Schematic of epsin1’s structure, which includes its ENTH domain (PDB: 1H0A) and its intrinsically disordered region. A schematic of clathrin recruitment to PI(4,5)P2 vesicles via epsin1 is included. **(B)** Representative fluorescent confocal images of vesicles that were tethered, thenincubated simultaneously with epsin1 and clathrin. Vesicles were fluorescently labeled with ATTO 488-DPPE. Epsin1 was fluorescently labeled with Alexa Fluor 594. Clathrin was fluorescently labeled with Alexa Fluor 647. Protein concentrations used in these images were 2 μM for epsin1 and 50 nM for clathrin. Vesicles used were composed of 73.5 mol% DOPC, 15 mol% DOPS, 10 mol% PI(4,5)P2, 1 mol% DPEG-15-biotin, and 0.5 mol% DPPE-ATTO488 and were extruded through 200 nm pores. Scale bars = 2 μm. Similar results were obtained across n = 3 independent replicates. **(C)** Histograms of vesicle diameters determined via confocal microscopy. n > 1,000 vesicles for each condition. For these fission experiments, vesicles, clathrin, and epsin1 were incubated at 37°C prior to tethering and imaging. Dashed lines represent the midline of the distribution for the control condition. **(D)** Summary of fission products from tethered vesicle experiments, expressed as the percent of vesicle diameters within the high fission product group of 50 nm or smaller. Error bars = mean ± first SD; n = 10 frames. For both **(C,D)** similar results were obtained across n = 3 independent replicates. Vesicles for **(C)** and **(D)** were composed of 73.5 mol% DOPC, 15 mol% DOPS, 10 mol% PI(4,5)P2, 1 mol% DPEG-15-biotin, and 0.5 mol% DPPE-ATTO647N and extruded through 200 nm pores.

Epsin1 alone induced membrane fission, producing smaller vesicle populations and a monotonic increase in fission products with increasing concentration (Fig. 6c-d). In contrast to clathrin’s inhibitory effect on amphiphysin1 mediated fission, the addition of clathrin enhanced epsin1-driven fission across the entire epsin1 concentration examined. Increasing the clathrin concentration from 25 nM to 50 nM further increased the extent of fission. Binding analyses show that clathrin increases the amount of epsin1 bound to vesicles, albeit to a lesser degree than amphiphysin1 (Fig. S14). These results indicate that clathrin assembles on epsin1 in a configuration such that its mechanical properties are favorable for inducing membrane fission.

## DISCUSSION

The principles uncovered here provide a framework for how clathrin lattices can regulate membrane fission. Notably, previous work has shown that enhancing clathrin assembly increases its curvature-sensing ability (*27*), whereas here we observe the opposite effect on curvature generation where enhanced clathrin assembly weakens membrane fission. While it is often thought that curvature sensing and generation are a manifestation of the same molecular behavior (*50*), our results present a system where this is not the case. Here, the mechanical state of the clathrin lattice determines whether clathrin acts as a stabilizer of pre-existing curvature or as a driver of new curvature, with these functions emerging in opposite mechanical regimes.

We observed a reciprocal relationship between membrane fission and lattice assembly across multiple perturbations; membrane fission was strongest when lattice assembly was weakened and suppressed when lattice assembly was enhanced, a trend that persisted even under conditions of high clathrin density on the membrane. These observations indicate that clathrin-mediated membrane remodeling is governed by lattice mechanics rather than protein abundance alone. The experimentally measured shifts in cage diameter provide a structural link between the assembly conditions and intrinsic lattice curvature. Consistent with this relationship, Brownian dynamics simulations identify triskelion geometry and lattice rigidity as independent determinants of curvature generation. Although lattice stiffness influenced curvature in the model, it was not measured experimentally and therefore remains a possible, but unconfirmed, contributor to the observed behavior.

Notably, fission increased as the intrinsic curvature of the clathrin cage increased (i.e., as cage diameter decreased), indicating that smaller, more highly curved lattices more effectively drove membrane fission. In our simulations, independently varying pucker angle and lattice stiffness altered lattice geometry and membrane bending capacity independent of binding kinetics, suggesting that the driving force for fission reflects the mismatch between the intrinsic curvature of the lattice and that of the underlying membrane. When the intrinsic cage diameter closely matched that of the starting vesicle population (∼200 nm), as observed in the pH 6.2 condition, this curvature mismatch was minimal and fission was negligible, consistent with reduced mechanical frustration. Thus, the experimentally observed relationship between smaller cage diameter and greater fission is consistent with a pucker-angle contribution to curvature mismatch. These observations suggest that modest environmental changes could bias clathrin toward membrane stabilization or deformation.

The influence of lattice geometry on membrane deformation may be particularly important during the late stages of fission, when the membrane neck narrows toward scission. Recent live-cell work proposed that clathrin polymerization around the base of Ω-shaped membrane profiles generates elastic forces that constrict and close the membrane pore (*22*). Although our experiments do not directly resolve pore constriction, and our simulations do not access near-scission geometries, our results provide a complementary physical framework for this model by showing that the membrane-deforming capacity of clathrin depends strongly on triskelion geometry and lattice stiffness. Thus, if clathrin acts at the pore as proposed, the efficiency of constriction may depend not simply on clathrin accumulation, but on the intrinsic curvature and mechanical state of the lattice assembled around the pore.

In addition to lattice rigidity, steric interactions among membrane-bound triskelia may also contribute to the mechanical response of the assembly. Under conditions favoring assembly, attractive protein-protein contacts incorporate triskelia into a cohesive lattice, reducing the population of unincorporated, membrane-bound components. When assembly is disfavored, this fraction of unincorporated, membrane-bound components increases, contributing to crowding and steric repulsion on the membrane (*17, 51*). The outward pressure generated by membrane crowding could aid curvature generation, particularly in regimes where clathrin is membrane bound but not yet fully assembled.

These principles extend to adaptor-mediated fission. Clathrin suppressed amphiphysin1-mediated fission yet enhanced epsin1-mediated fission, indicating that different adaptors bias clathrin into mechanically distinct lattice configurations. Given that the presence of clathrin increased amphiphysin1 binding, we suspect that clathrin assembles on top of amphiphysin1 in such a way that its mechanical properties are inhibiting the ability of amphiphysin1’s BAR domain and intrinsically disordered region (IDR) to drive membrane curvature. Within epsin1, however, interactions between the IDR and clathrin may elicit lattice mechanical properties that work cooperatively with epsin1’s innate ability to drive fission. Given that epsin1 is an early-stage adaptor protein (*13*), it is possible that these interactions are essential for promoting membrane bending during the beginning of endocytosis. In contrast, these interactions may be less essential in amphiphysin1, which is a mid-to late-stage adaptor protein (*52*). Thus, adaptor proteins may not merely recruit clathrin to membranes but actively tune its mechanical contribution to curvature generation, stability, and fission.

By identifying intrinsic lattice curvature and organization, rather than clathrin abundance alone, as key determinants of fission, this work suggests a revised view of clathrin’s function in endocytosis in which clathrin alternates between curvature generating and stabilizing mechanical states that collectively shape the pathway’s timing, efficiency, and responsiveness to cellular context. Because many membrane-deforming processes rely on the assembly of intrinsically curved protein scaffolds, including caveolin, COPI/II coats, and ESCRT assemblies (*53, 54*), similar shifts in assembly-dependent mechanical state may broadly determine whether protein assemblies stabilize membrane curvature or actively drive fission.

## MATERIALS AND METHODS

### Materials

1,2-dioleoyl-sn-glycero-3-phosphocholine (DOPC), 1,2-dioleoyl-sn-glycero-3-phospho-L-serine (sodium salt) (DOPS), 1,2-dioleoyl-sn-glycero-3-[(N-(5-amino-1-carboxypentyl) iminodiacetic acid)succinyl] nickel salt (DGS-NTA-Ni^2+^), and L-α-phosphatidylinositol-4,5-bisphosphate ammonium salt (PI-(4,5)-P2) were purchased from Avanti Polar Lipids. Dipalmitoyl-decaethylene glycol-biotin (DP-EG15-biotin) was generously provided by D. Sasaki from Sandia National Laboratories, Livermore, CA(*55*). ATTO-488 1,2-Dipalmitoyl-sn-glycero-3-phosphoethanolamine (DPPE), ATTO 647N 1,2-Dipalmitoyl-sn-glycero-3 phosphoethanolamine (DPPE), and ATTO-488 maleimide were purchased from ATTO-TEC. NeutrAvidin, Alexa Fluor 594 NHS Ester, Alexa Fluor 647 NHS Ester, Pierce HRV 3C Protease Solution Kit, β-mercaptoethanol (BME), sodium chloride, 4-(2-hydroxyethyl)-1-piparazineethanesulphonic acid (HEPES), tris(hydroxymethyl)aminomethane (Tris), Triton X-100, Zeba spin desalting columns (7k MWCO, 0.5 mL), Gibco penicillin-streptomycin-glutamine (100x), and Gibco fetal bovine were purchased from Thermo Fisher Scientific. Isopropyl-beta-D-thiogalactoside (IPTG) and dithiothreitol (DTT) were purchased from Gold Biotechnology. Phenylmethanesulfonyl fluoride (PMSF), imidazole, poly-L-lysine (PLL), Thrombin CleanCleave Kit, ammonium sulfate, calcium chloride, and tissue-culture treated culture dishes were purchased from Sigma-Aldrich. HyClone Trypsin Protease, 0.05%, HyClone DMEM high glucose without L-glutamine, sodium pyruvate, HyClone DMEM high glucose modified without phenol red, sodium pyruvate, L-glutamine, and HyClone Ham’s Nutrient Mixture F12 were purchased from Cytiva Life Sciences.

### Protein expression and purification

### Clathrin

Clathrin heavy chain (CHC) and clathrin light chain (CLC) were expressed and purified as described below utilizing constructs described previously(*27*). The proteins were stored separately at-80°C and immediately before use were mixed in a 1:1 molar ratio.

### Clathrin Heavy Chain (CHC)

BL21 competent cells (NEB) were transformed with CHC expression plasmid pET28A (+) 6His-ratCHC-FL, grown in 2xTY + 1/20 volume of 10xM9 salts, 50 µg/ml kanamycin, pH 7.4 at 30°C to an OD600 of 1.4, cooled to 12°C, induced with 1 mM IPTG, and expressed for 24 hr at 12°C. Pellets were frozen and stored at-80°C. For purification of CHC, pellets from 2 L of cultured cells were defrosted on ice, suspended in 80 mL of 0.5 M Tris-HCl, pH 8.0, 10 mM BME, 1% Triton X-100, 3 tablets of EDTA-free protease inhibitor cocktail (Roche), and sonicated at 4 x 60% amplitude on ice. The cells/lysate was then spun down in an ultracentrifuge at 103,800 x g for 30 minutes at 4°C. The lysate was then batch absorbed to 10 mL bed volume Ni-NTA agarose resin (Millipore Sigma) for at minimum 1 hour at 4°C with stirring at 60 rpm. The column was then washed with (1) 5 column volumes 0.4 M Tris-HCl pH 8.0, 10 mM imidazole, 1 mM PMSF, 10 mM BME, (2) 5 column volumes 0.4 M Tris-HCl pH 8.0, 20 mM imidazole, 1 mM PMSF, 10 mM BME, (3) 5 column volumes 0.4 M Tris-HCl and 1 M NaCl pH 8.0, 20 mM imidazole, 1 mM PMSF, 10 mM BME, (4) 5 column volumes 0.4 M Tris-HCl pH 8.0, 20 mM imidazole, 1 mM PMSF, 10 mM BME and eluted slowly with 5 column volumes of 0.5 M Tris-HCl pH 8.0, 200 mM imidazole, 1 mM PMSF, 10 mM BME. The eluate was then concentrated with a 30kDa MWCO Amicon Ultra – 15 centrifugal concentrator (Millipore) and spun at 4,000 x g for one hour. The concentrated eluate was then run on a Superdex 200 HiLoad chromatography column (Cytiva) at 4°C. The protein fractions containing CHC were pooled, concentrated with a 30kDa MWCO Amicon Ultra-4 centrifugal concentrator (Millipore) to a final concentration of 15-20 µM, and dialyzed overnight into 0.5 M Tris-HCl, pH 8.0, 1 mM EDTA, 5 mM DTT. Purified CHC was aliquoted in 0.6 mL microcentrifuge tubes (Eppendorf), flash frozen in liquid nitrogen, and stored at-80°C.

### Clathrin Light Chain (CLC)

CLC was expressed together with CHC, and the CHC was removed by heat denaturation as follows: BL21 competent cells (NEB) were transformed with CHC expression plasmid pET28A (+) 6His-ratCHC-FL and pBAT4-no tag CLCA1 (co-expressed with pET28A (+) 6His-ratCHC-FL), and the expression and purification were carried out as described above with the addition of 100 µg/ml ampicillin to select for the CLC plasmid, until the concentration of the affinity chromatography eluate. The CLC eluate was concentrated by precipitation with an equal volume of saturated ammonium sulfate (in 0.5 M Tris-HCl, pH 7.0), stirred at 60 rpm for 30 minutes at 4°C, collected by centrifugation at 18,000 x g for 40 minutes at 4°C, and the pellet was dissolved in 0.5 M Tris-HCl pH8.0, 1 mM EDTA, 5 mM DTT. The resuspended pellet was then heated in a water bath at 90°C for 5 minutes to denature and precipitate the CHC protein which was removed by centrifugation at 8,600 x g for 5 minutes at room temperature. The sample was then applied to a Superdex 75 10/300 GL chromatography column (Cytiva) at 4°C. The protein fractions containing CLC were pooled, concentrated with a 10 kDa MWCO Amicon Ultra – 4 centrifugal concentrator (Millipore) at 3,000 x g for 30 minutes, and CLC was dialyzed against 0.5 M Tris-HCl, pH 8.0, 1 mM EDTA, 5 mM DTT at 4°C overnight. The purified CLC was aliquoted in 0.6 mL microcentrifuge tubes (Eppendorf), flash frozen in liquid nitrogen, and stored at-80°C.

### Amphiphysin1

Amphiphysin1 was expressed as a fusion protein containing N-terminal glutathione-S-transferase (GST) in BL21 (DE3) cells. BL21 cells were transformed with amphiphysin1 expression plasmid pGex4T2, grown in 2xTY pH 7.4, 100 µg/mL ampicillin, at 37°C to an OD600 of 0.6, cooled to 30°C, induced with 1 mM IPTG, and expressed for 6 hours. Pellets were frozen and stored at-80°C. For purification, pellets from 2 L of culture were thawed on ice, suspended in 40 mL of 0.5 M Tris-HCl, 5 mM EDTA, 5% v/v glycerol, 10 mM BME, 1 mM PMSF, 1% v/v Triton X-100, and 2 tablets of EDTA-free protease inhibitor cocktail (Roche), and sonicated at 4 x 60% amplitude on ice. The cells/lysate was then spun down in an ultracentrifuge at 103,800 x g for 30 minutes at 4°C. The lysate was then batch absorbed to 2 mL bed volume glutathione agarose resin (Thermo Fisher) for at minimum 1 hour at 4°C with stirring at 60rpm. The column was then washed with (1) 10x column volume of 0.4 M Tris-HCl, 1 M NaCl, 10 mM BME, 1 mM PMSF pH 8.0, (2) 10x column volume of 25 mM HEPES, 150 mM NaCl, 1 mM EDTA, 10 mM BME pH 7.4 and eluted with 4 column volumes of 150 mM NaCl, 50 mM Tris, 20 mM GSH, 1 mM EDTA pH 8.0. The eluate was concentrated with a 30kDa MWCO Amicon Ultra – 15 centrifugal concentrator (Millipore) and incubated with soluble HRV-3C protease overnight at 4°C. Excess HRV-3C, which contained a GST tag, was removed by passage through another glutathione agarose column. Amphiphysin1 was then exchanged into 25 mM HEPES, 150 mM NaCl, and 5 mM DTT pH 7.4 and the purified protein was aliquoted, flash frozen, and stored at-80°C.

### Epsin1

Epsin1 was expressed as a fusion protein containing N-terminal glutathione-S-transferase (GST) in BL21 (DE3) cells. BL21 cells were transformed with epsin1 expression plasmid pGex6P1 and the expression and purification were carried out as described above for amphiphsyin1, until the glutathione resin elution. Epsin1 was cleaved directly from the resin by combining with thrombin and allowed to mix overnight at 4°C. The cleaved protein was collected and combined with p-aminobenzamidine agarose (Thermo Fisher) to remove thrombin for 1 hour at 4°C. The protein was then aliquoted and stored at-80C.

His-ENTHΔH0: BL21 (D32) bacterial cells were transformed with his-ENTHΔH0 expression plasmid pRSET-his-ENTHΔH0 grown in 2xTY + 100 µg/ml ampicillin, pH 7.4 at 37°C to an OD600 of 0.6, cooled to 18°C, induced with 1 mM IPTG, and expressed overnight at 18°C. Pellets were frozen and stored at-80°C. Pellets from 2 L of cultured cells were defrosted on ice, suspended in 40 mL of 0.5 M Tris-HCl, pH 8.0, 10 mM BME, 5% v/v glycerol, 1 mM PMSF, 4 tablets of EDTA-free protease inhibitor cocktail (Roche), and sonicated at 4 x 60% amplitude on ice. The cells/lysate was then spun down in an ultracentrifuge at 103,800 x g for 30 minutes at 4°C. The lysate was then batch absorbed to 3 mL bed volume Ni-NTA agarose resin (Millipore Sigma) for at minimum 1 hour at 4°C with stirring at 60 rpm. The column was then washed with (1) 5 column volumes 0.4 M Tris-HCl pH 8.0, 10 mM imidazole, 1 mM PMSF, 10 mM BME, (2) 5 column volumes 0.4 M Tris-HCl pH 8.0, 20 mM imidazole, 1 mM PMSF, 10 mM BME, (3) 5 column volumes 0.4 M Tris-HCl and 1 M NaCl pH 8.0, 20 mM imidazole, 1 mM PMSF, 10 mM BME, (4) 5 column volumes 0.4 M Tris-HCl pH 8.0, 20 mM imidazole, 1 mM PMSF, 10 mM BME and eluted slowly with 5 column volumes of 0.5 M Tris-HCl pH 8.0, 200 mM imidazole, 1 mM PMSF, 10 mM BME. Peak fractions were pooled, concentrated, and buffer exchanged into 25mM HEPES, 150 mM NaCl (pH 7.4) 5 mM DTT and aliquoted, flash frozen, and stored at - 80°C.

### Protein Labeling

ATTO-488 NHS-ester, Alexa Fluor 594 NHS-ester, and Alexa Fluor 647 NHS-ester were dissolved in dimethyl sulfoxide (DMSO) at concentrations of 5 mM and 10 mM (ATTO-488), 10 mM (Alexa Fluor 594) and 5 mM (Alexa Fluor 647) and stored at-80°C.

### Clathrin

Clathrin was freshly labeled prior to each experiment. Clathrin light chain and clathrin heavy chain were thawed on ice and combined in a 1:1 stoichiometric ratio to yield a solution of 2-3 µM triskelia. Using a 7 kDa MWCO Zeba spin column (Thermo Fisher), the clathrin was exchanged into 100 mM sodium bicarbonate and 20 mM BME (pH 8.2). The dye solution was added to the clathrin solution at a stoichiometric ratio of 6 dyes per triskelion. This mixture reacted for 20 minutes at room temperature and was then immediately exchanged into a storage buffer containing 10 mM Tris-HCl and 20 mM BME (pH 8.0). Clathrin was then centrifuged for 10 minutes at 9,000 x g to remove any aggregates. The final concentration of clathrin was 1-1.5 µM and the labeling ratios varied from 0.5 to 1 dye per triskelion. Protein and dye concentrations were measured using UV-Vis spectroscopy.

### Amphiphysin1, Epsin1, and his-ENTHΔH0

Primary amines within these proteins were labeled in buffer consisting of 25 mM HEPES, 150 mM NaCl, and 20 mM BME (pH 7.4). Protein concentration varied from 25-100 µM. The dye solution was added to 100 µL of protein such that the DMSO never exceeded 1 v/v% and the stoichiometric ratio of dye to protein was 2 to 1. The mixture was then allowed to react for 30 minutes at room temperature. Unreacted dye was removed using Zeba spin columns (Thermo Fisher) and protein and dye concentrations were measured using UV-Vis spectroscopy. The resulting labeling ratios varied from 0.8 to 1.2 dyes per protein and labeled proteins were stored as 5 µL aliquots at-80°C.

### Vesicle Preparation

Lipid aliquots were thawed from-80°C and combined in the following molar percentages for use in the tethered vesicle assays:

- 88.5% DOPC, 10% DGS-NTA-Ni^2+^, 1% DP-EG15-biotin, 0.5% DPPE-ATTO-647N
- 78.5% DOPC, 20% DGS-NTA-Ni^2+^, 1% DP-EG15-biotin, 0.5% DPPE-ATTO-647N
- 78.5% DOPC, 15% DOPS, 5% Brain PI(4,5)P2, 1% DP-EG15-biotin, 0.5% DPPEATTO-647N
- 73.5% DOPC, 15% DOPS, 10% Brain PI(4.5)P2, 1% DP-EG15-biotin, 0.5% DPPEATTO-647N
- 78.5% DOPC, 15% DOPS, 5% Brain PI(4,5)P2, 1% DP-EG15-biotin, 0.5% DPPEATTO-488
- 73.5% DOPC, 15% DOPS, 10% Brain PI(4.5)P2, 1% DP-EG15-biotin, DPPE-0.5% ATTO-488

For TEM:

- 90% DOPC, 10% DGS-NTA-Ni^2+^

Solvents were evaporated with a nitrogen stream, followed by vacuum storage for at least 2 hours. The lipid film was then hydrated to a lipid concentration of 500 µM in buffer containing 25 mM HEPES, 150 mM NaCl, and 20 mM BME (pH 7.4) For vesicles containing PI(4,5)P2, 0.5 mM EDTA and 0.5 mM EGTA were included in the buffer to prevent aggregation of PI(4,5)P2 by divalent metal contamination. Vesicles containing PI(4,5)P2 were also prepared fresh each day of use due to the relatively quick degradation of brain PI(4,5)P2 when exposed to oxygen(*56*). Rehydrated lipid suspensions were thoroughly mixed and held at room temperature for 15 minutes and subjected to 5 freeze/thaw cycles by submerging the solution in liquid nitrogen for 1 minute and thawing at 50°C. The vesicles were then extruded through a 200 nm polycarbonate membrane (Whatman).

### Tethered Vesicle Assay

PLL-PEG was synthesized as described previously(*57, 58*). Imaging wells consisted of 1.6 mm thick silicone gaskets (Grace Bio-Labs) with 5 mm diameter circular holes cut into them. These gaskets were placed directly on top of no. 1.5 glass coverslips. Prior to use, gaskets and coverslips were thoroughly cleaned using Hellmanex (Hellma Analytics) and dried under nitrogen. Once assembled, 30 µL of PLL-PEG was added and allowed to incubate for 20 minutes. The PLL-PEG was rinsed out with 25 mM HEPES, 150 mM NaCl (pH 7.4) and 6 µg of NeutrAvidin dissolved in 25 mM HEPES, 150 mM NaCl (pH 7.4) was added to each sample well and allowed to incubate for 10 minutes. Wells were then rinsed with the buffer of the solution being tethered to remove excess NeutrAvidin. Vesicles and proteins were then tethered for 5-10 minutes before being rinsed to remove any untethered vesicles. Slides were imaged immediately.

### Fission Assay

Fission experiments with clathrin only were performed by mixing DGS-Ni-NTA containing vesicles (molar percentages above) at a concentration of 10 µM with unlabeled protein at specified concentrations in 25 mM HEPES, 150 mM NaCl, 20 mM BME at pH 6.2, 7.4, and 8.3. For conditions with calcium, the buffer 25 mM HEPES, 150 mM NaCl, 1 µM CaCl_2_, 20 mM BME (pH 7.4) was used. The vesicles and protein were then incubated at 37°C for 30 minutes. After incubation, the mixture was added to the wells and allowed to tether for 10 minutes prior to being rinsed to remove untethered vesicles.

Fission experiments with adaptor proteins amphiphysin1 and epsin1 were performed by mixing PI(4,5)P2 vesicles (78.5% DOPC, 15% DOPS, 5% Brain PI(4,5)P2, 1% DP-EG15-biotin, 0.5% ATTO-647N for amphiphysin1, 73.5% DOPC, 15% DOPS, 10% Brain PI(4.5)P2, 1% DP-EG15-biotin, 0.5% ATTO-647N for epsin1) at a concentration of 10 µM with unlabeled proteins at specified concentrations in 20 mM HEPES, 150 mM NaCl, 20 mM BME, 0.5 mM EDTA, 0.5 mM EGTA (pH 7.4). To ensure that the adaptor proteins did not interact with the vesicles before clathrin was introduced, clathrin was added before amphiphysin1 or epsin1. The mixture was then incubated and tethered as described above. To determine binding stoichiometries, ATTO-647N was replaced with 0.5% ATTO-488, amphiphysin1 and epsin1 were labeled with Alexa-594 as described above, and clathrin was labeled with Alexa-647 as described above. After tethering, vesicles were allowed to incubate 30 minutes prior to imaging and wells were not rinsed.

### Binding Curves

To produce binding curves, vesicles composed of 88.5% DOPC, 10% DGS-NTA-Ni, 1% DP-EG15-biotin, 0.5% ATTO-647N were tethered to a glass coverslip and sufficiently rinsed to remove vesicles that were not tethered. Protein labeled with ATTO-488 was then added to the wells at the specified concentrations and allowed to incubate for 30 minutes prior to imaging.

### Fluorescent Microscopy for Tethered Vesicle Assay

Imaging was performed on a laser scanning confocal microscope (Leica Stellaris 5). Three excitation lasers were used: 488 nm, 561 nm, and 638 nm. A Leica, HC PL APO 63x, 1.4 NA oil-immersion objective was used to acquire images, and the zoom factor was set such that square pixels had dimensions of 70 nm x 70 nm. All images were acquired using a scan speed of 600 Hz.

### Image Processing for Tethered Vesicle Assay

Image processing was performed similarly to previous work(*57*). Images of diffraction limited, fluorescent lipid and protein puncta were acquired using confocal microscopy. Each image in each channel (up to three channels) was a three-frame repeat of a single field of view. The fluorescent amplitudes of all diffraction limited puncta were obtained using publicly available cmeAnalysis particle detection software and vesicles were detected by fitting 2D Gaussian profiles to each puncta(*59*). The first frame of the lipid fluorescence channel served as our reference, and to be included in analysis puncta had to persist in the same location through three consecutive frames and have amplitudes significantly higher than the fluorescent background. Protein binding for each vesicle was determined by analyzing the colocalization between the lipid channel and protein channel(s). The search radius for colocalized protein puncta was equal to 3 times the standard deviation of the Gaussian fit of the lipid channel.

### Calibration of vesicle diameters and number of bound proteins

Vesicle diameters and the number of proteins bound to each vesicle were determined as previously described(*26*). Briefly, tethered vesicles were imaged to obtain a distribution of the fluorescence intensities for a protein-free vesicle population. A conversion factor from fluorescence intensity to size was determined by overlaying the square root of the fluorescence distribution and the dynamic light scattering distribution. (Fig. S1) The number of proteins bound to each vesicle was determined using single-molecule imaging to determine the brightness of a single ATTO-488, Alexa-594, and Alexa-647 labeled protein. (Fig. S2). The fluorescent proteins were added to clean glass coverslips at 50 pM and the intensities of the punctate structures were used to generate a distribution. The peak value of this fluorescence intensity distribution represented the intensity of a single fluorescently labeled protein. Quantitative adjustments were made during data processing to account for changes in image frame accumulates and linear gain settings on the detector.

The data in Figures 2B and 3B were fit using a Langmuir-Hill adsorption isotherm to determine if the binding of his-clathrin to the membrane is a cooperative process, as described previously(*27*) (Eq. 1). Here, “n” corresponds to the Hill coefficient.

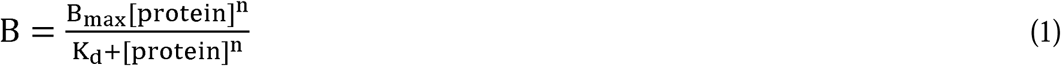

### Transmission Electron Microscopy

Vesicles for TEM were composed of 90% DOPC, 10% DGS-NTA-Ni. Dried lipid films were rehydrated in 25 mM HEPES, 150 mM NaCl (pH 7.4) and extruded through a 200nm filter as described above. Fission experiments were performed as described above: vesicles were incubated at 10 µM with clathrin diluted to the indicated concentration at 37°C for 30 minutes. For protein-free vesicle samples, vesicles were diluted to 100 µM. For each condition, 5 µL of sample was placed onto a glow-discharged, 200 square mesh, formvar supported carbon-coated grid (Electron Microscopy Sciences) for one minute. Remaining liquid was wicked away with filter paper, and the grid was stained with 1% uranyl acetate for 45 seconds twice. Images were collected on a Talos F200C G1 Transmission Electron Microscope and analyzed using ImageJ software.

### Membrane Bending Simulations

Meso-scale simulations of membrane bending by clathrin were carried out using a continuum membrane model coupled to rigid-body coarse-grained (CG) clathrin triskelia joined by harmonic potentials, as elaborated in our recent work(*34*). The dynamic membrane itself is modeled by the method presented by Lin and Brown(*35*). Briefly, the lipid bilayer free energy is modeled according to the Helfrich functional(*60*) in the small gradient approximation,

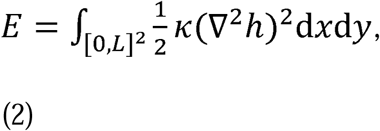

where κ is the membrane bending modulus (20 k_B_T) and *L* = 200nm is the dimension of the square base plane over which the membrane is represented with periodic boundary conditions (PBC). *h=h(x,y)* is the height of the membrane above the base plane (Monge gauge)(*61*).The stochastic membrane dynamics are propagated in Fourier space consistent with low Reynolds number hydrodynamics:

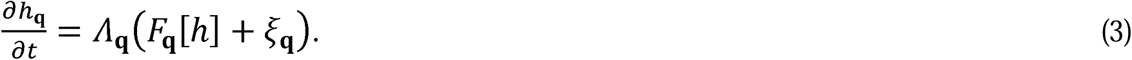

Here **q** is the Fourier mode wave vector, ⋀_**q**_ 1/4*ηq* with viscosity *η =* 24 k_B_ T ns nm^-3^, ξ**_q_** is a Gaussian noise satisfying the fluctuation-dissipation relation, and *F_q_* is the Fourier transform of the force per unit area in the z-direction on the membrane,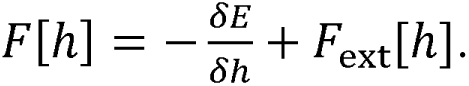 This last term, *F _ext_* accounts for external forces on the membrane, through which we couple our dynamic membrane to our CG triskelia via harmonic springs. We use a grid size of 55 in both x and y for a total of 3025 independent Fourier modes.

The CG clathrin model consists of individual triskelia treated as rigid bodies with three binding sites for other triskelia (Fig. S11). A flexible coat is then created by imposing harmonic potentials on the length and angles of bonds between the clathrin-binding sites. Each clathrin-clathrin bond is parametrized by 4 coordinates: the bond length σ, bond angles θ_1_ and θ_2_, and dihedral angle ω (see Fig. S11). We assign a harmonic energy 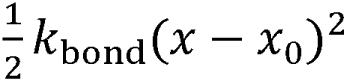 to each of these coordinates, with respective rest values of σ = 5nm, θ_1_ = θ_2_ = π, and ω = 0. We use the same numerical k_bond_ value for all parameters as we have only one meso-scale elastic parameter against which to calibrate: its effective bending modulus κ_clat._ This is the most widely studied rigidity parameter for clathrin coats and reported values vary from as little as 20 k_B_T to in excess of 500 k_B_T(*39, 40, 62–64*). Our range of k_bond_ corresponds to a range of κ_clat_values roughly 10 k_B_T to 200 k_B_T, as measured via analysis of the stress-strain relation of clathrin buckling simulations(*65, 66*), covering much of the proposed rigidity space, from below membrane rigidity to an order of magnitude larger. Individual triskelia centers of mass are connected to the membrane via harmonic potentials in the z-direction with a rest length of 15 nm and a spring constant of 1000 k_B_T/nm^2^. We use HOOMD-blue to propagate the triskelia with a Brownian integrator(*67*), with spring coupling to the membrane implemented as an external force on each triskelion, propagating the membrane and triskelia together consistently. The translational and rotational friction constants for the integrator are chosen to reproduce the triskelion diffusion constants used in previous work(*33*). The simulation timestep is Δ*t* 0.01 ns.

In order to approximate the stochastic growth of the clathrin lattice in our simulations, we attempt to add a new triskelion at fixed time intervals (10 μs, 50 μs, and 150 μs were used in different simulation series, Fig. S11) starting from an initial pre-formed state of six triskelia bonded together in a loop, adding a maximum of 16 additional triskelia (22 total). While these binding kinetics are not constrained to a rigorous equation of motion(*68*), they allow us to test how assigning a fast characteristic rate to clathrin lattice growth will impact the dynamics of the membrane remodeling. We do not allow clathrin triskelia to dissociate from the lattice or membrane. With this approximation, we cannot assess the probability that clathrin lattices will nucleate and grow(*33*), and thus we do not comment on that with the model. However, given that the experiments are at a high enough concentration to drive nucleation and growth of lattices at all conditions tested, our model can then rigorously compare how lattice mechanics will impact the degree of membrane invagination driven by assembled lattices. Each triskelion addition is made at the closest unoccupied clathrin binding site to the center of the initial hexamer in order to create approximately circular coats. If the newly added triskelion would have its center of mass within 14 nm of an already present triskelion, the addition is canceled and the next closest binding site is chosen. Additionally, if at least 12 triskelia have already been added and the cage curvature already exceeds 0.015 nm^-1^ (radius 66 nm), triskelion addition is canceled in order to prevent the cage from driving our membrane model beyond what is representable in Monge gauge. At 25 evenly spaced time points between each triskelion addition, we check for open clathrin binding sites within 7 nm of one another, irreversibly bonding together any such pairs found. The membrane was allowed to equilibrate for 100 μs (longer than the relaxation time of the slowest Fourier mode) before the first triskelion addition. The final triskelion addition was followed by another 100 μs equilibration period, the latter 50 μs of which was used to calculate the equilibrium measurements presented in Fig. 4b. Clathrin lattice curvature was measured via least-squares fitting of all triskelion centers of mass to the fourth-order Taylor expansion of a hemisphere.

## ACKNOWLEDGEMENTS

Thank you to Eileen M. Lafer and Liping Wang for generously providing the pET28A (+) CHC and pBAT4 CLC plasmids, as well as troubleshooting assistance with CHC and CLC purification protocols. TEM images were acquired at the Core Center of Excellence in Nano Imaging at the University of Southern California with aid from Carolyn B. Marks. S.L.F. gratefully acknowledges postdoctoral funding support from the Gordon and Betty Moore Foundation. This research was supported by the National Institutes of Health through the following grants: R35GM147333 to N.B. and W.F.Z; R35GM133644 to S.L.F and M.E.J.

