## Supporting Information for "Clathrin is an Intrinsic Driver of Membrane Fission"

### Supplementary Figures

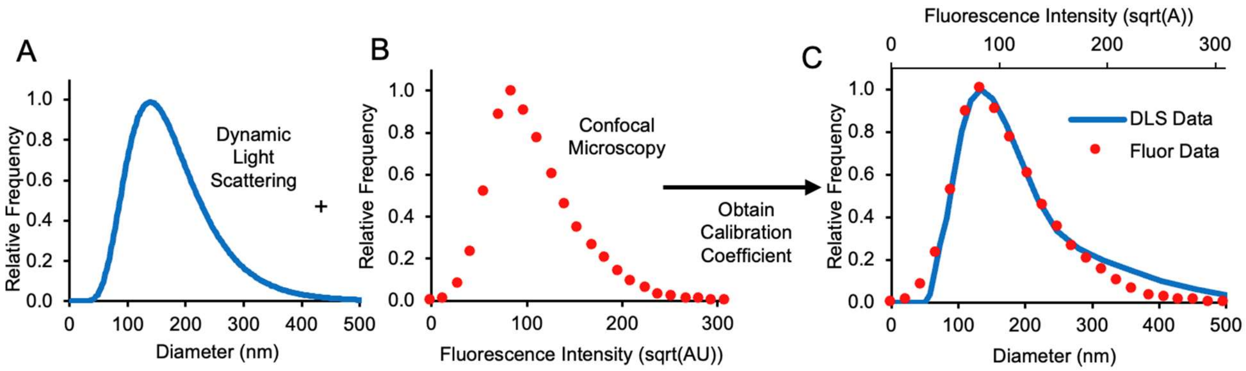


**Figure S1 – Calibration of Vesicle Diameters vis Dynamic Light Scattering**

(**A**) Representative DLS histogram for vesicles that were extruded through 200 nm pore membranes. **(B)** Distribution of fluorescent intensities for the same vesicles from panel **A** that were tethered to glass coverslips and imaged via fluorescence microscopy. The square roots of fluorescence intensities were used since fluorescence intensity scales linearly with vesicle surface area (i.e. intensity ~ $D^2$). **(C)** Overlay of histograms for the extruded vesicles measured by DLS in **A** and fluorescence intensity distribution in **B**. This comparison was obtained by using a scaling factor that was derived from the ratio of the distribution values in **A** and **B**.

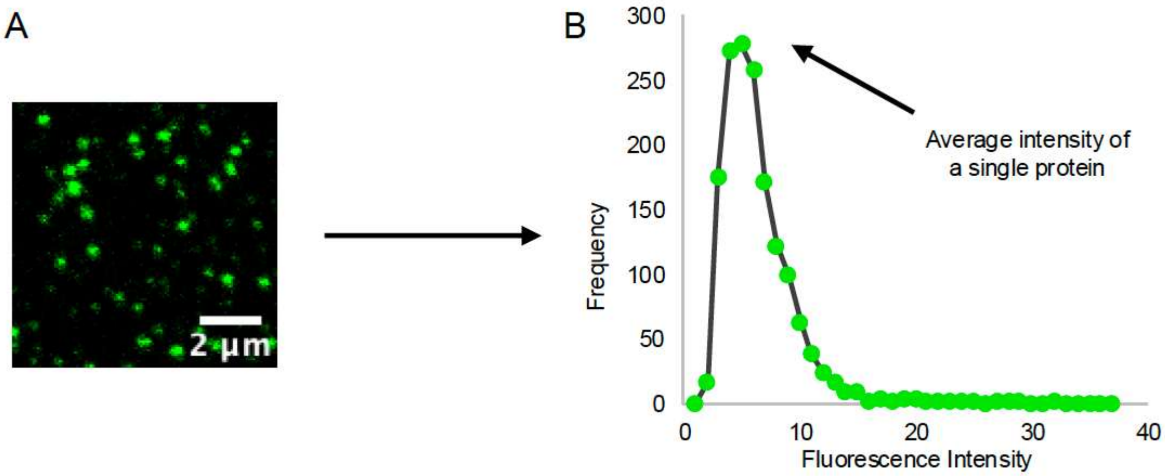


**Figure S2 – Single molecule imaging to calibrate protein copy number**

(**A**) Representative image of ATTO-488 labeled his-clathrin that was absorbed to a glass coverslip at 50 pM. **(B)** Representative fluorescent intensity distribution for imaged puncta.

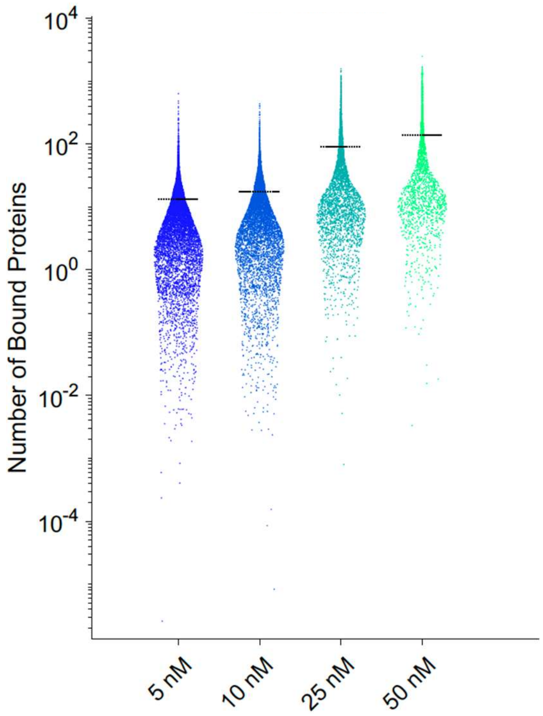


**Figure S3 – Number of bound clathrin triskelia per vesicle after fission and tethering**

The raw data for the average binding depicted in Figure 1F. Vesicles were composed of 88.5% DOPC, 10% DGS-NTA-NI$^{2+}$, 1% DPEG-15-biotin, and 0.5% ATTO-647N, and extruded through 200 nm pores. Each dot represents a vesicle, and the dashed line represents the mean of the data.

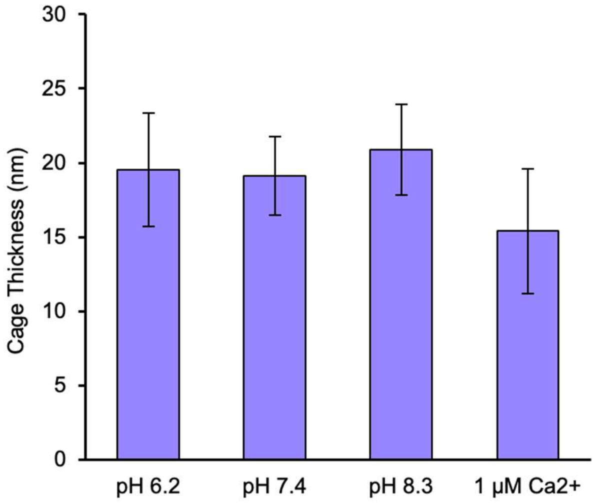


**Figure S4 – Measured thickness of clathrin cages.**

The mean thickness of clathrin cages assembled on 90% DOPC, 10% DGS-NTA, 200 nm extruded vesicles measured from negative stain TEM micrographs. Clathrin concentration for each condition was 50 nM. Error bars represent 1 SD; pH 6.2 n = 21, pH 7.4 n = 19, pH 8.3 n = 22, 1 µM $Ca^{2+}$ n = 18.

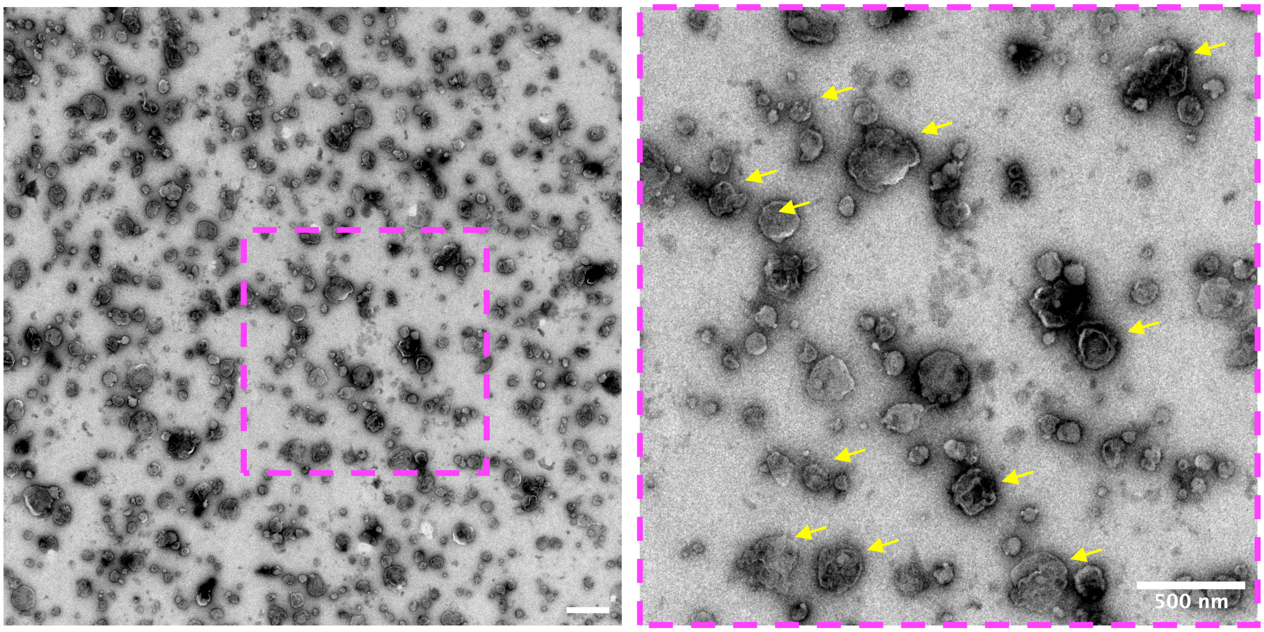


**Figure S5 – TEM sample preparation causes substantial vesicle rupture in the absence of clathrin.**

Micrograph of 90% DOPC and 10% DGS-NTA vesicles affixed to EM grid with 1% uranyl acetate. Many vesicles (examples denoted with arrows) ruptured during the staining process.

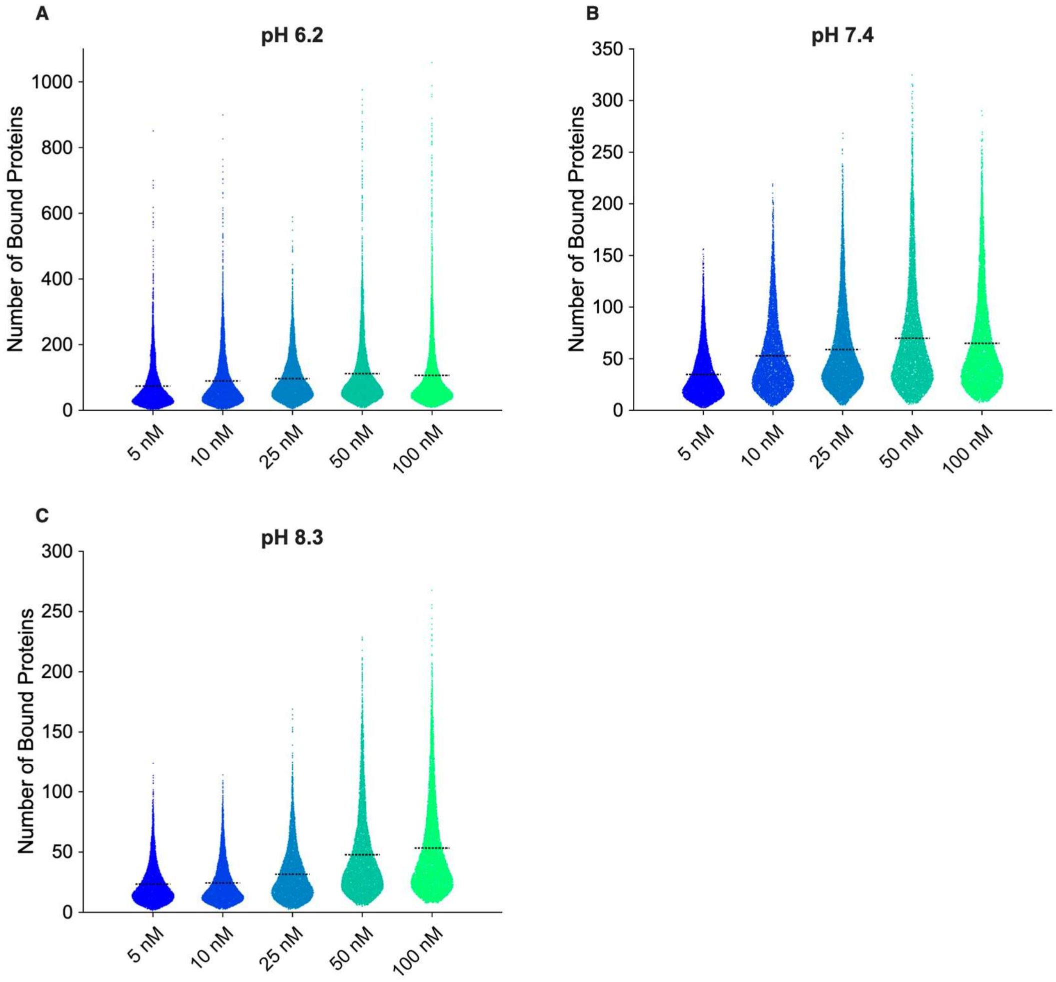


**Figure S6 – Number of bound clathrin triskelia per vesicle as a function of pH**

The raw data for the binding curves depicted in Figure 2B. Vesicles were composed of 88.5% DOPC, 10% DGS-NTA-NI$^{2+}$, 1% DPEG-15-biotin, and 0.5% ATTO-647N, and extruded through 200 nm pores. Each dot represents a vesicle for increasing concentrations of clathrin for **(A)** pH 6.2 **(B)** pH 7.4 and **(C)** pH 8.3. The dashed line represents the mean.

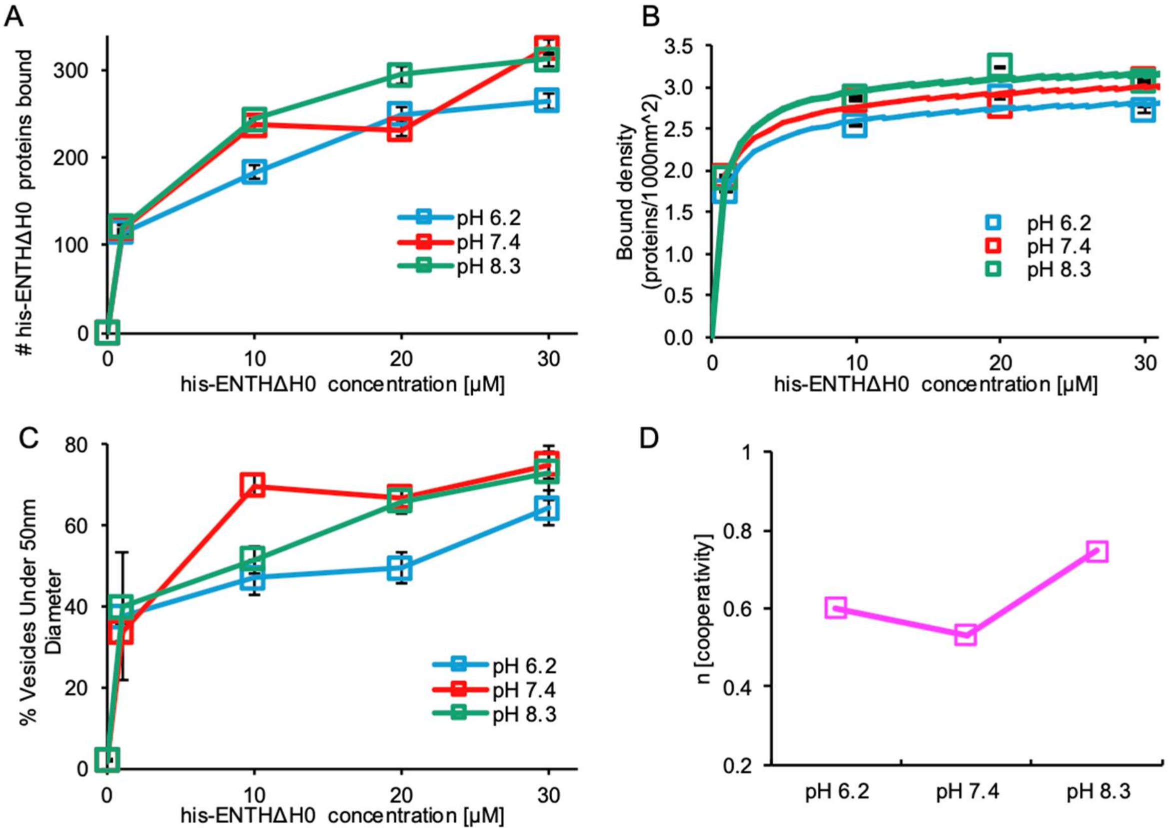


**Figure S7 – Bulk pH does not alter membrane binding and fission by his-ENTHΔH0**

**(A)** Number of his-ENTHΔH0 ATTO-488 labeled proteins bound to vesicles at pH 6.2, 7.4, and 8.3. **(B)** Binding isotherms and their corresponding lines of best fit for his-ENTHΔH0. This data was fit using Supporting Equation S1. **(C)** Percent of vesicles below 50 nm in diameter after incubation with his-ENTHΔH0 at increasing concentrations **(D)** Regressed values of n for the binding isotherms of his-ENTHΔH0. Vesicle composition for **(A-D)**: 78.5% DOPC, 20% DGS-NTA, 1% DPEG-15-biotin, 0.5% ATTO-647N, extruded through 200 nm pores.

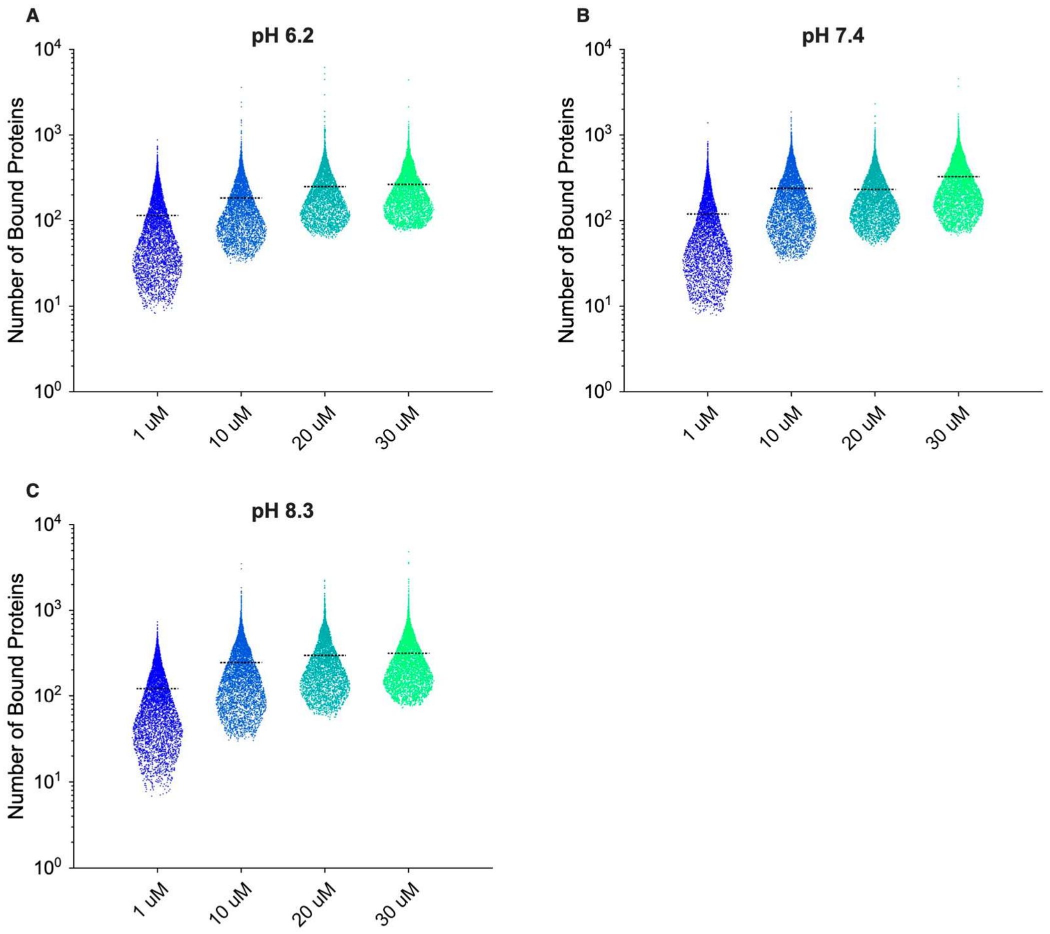


**Figure S8 – Number of bound his-ENTHΔH0 proteins per vesicle as a function of pH**

The raw data for the binding curves depicted in Figure S7. Vesicles were composed of 78.5% DOPC, 20% DGS-NTA-$NI^{2+}$, 1% DPEG-15-biotin, and 0.5% ATTO-647N and extruded through 200 nm pores. Each dot represents a vesicle for increasing concentrations of his-ENTHΔH0 for **(A)** pH 6.2 **(B)** pH 7.4 and **(C)** pH 8.3. The dashed line represents the mean.

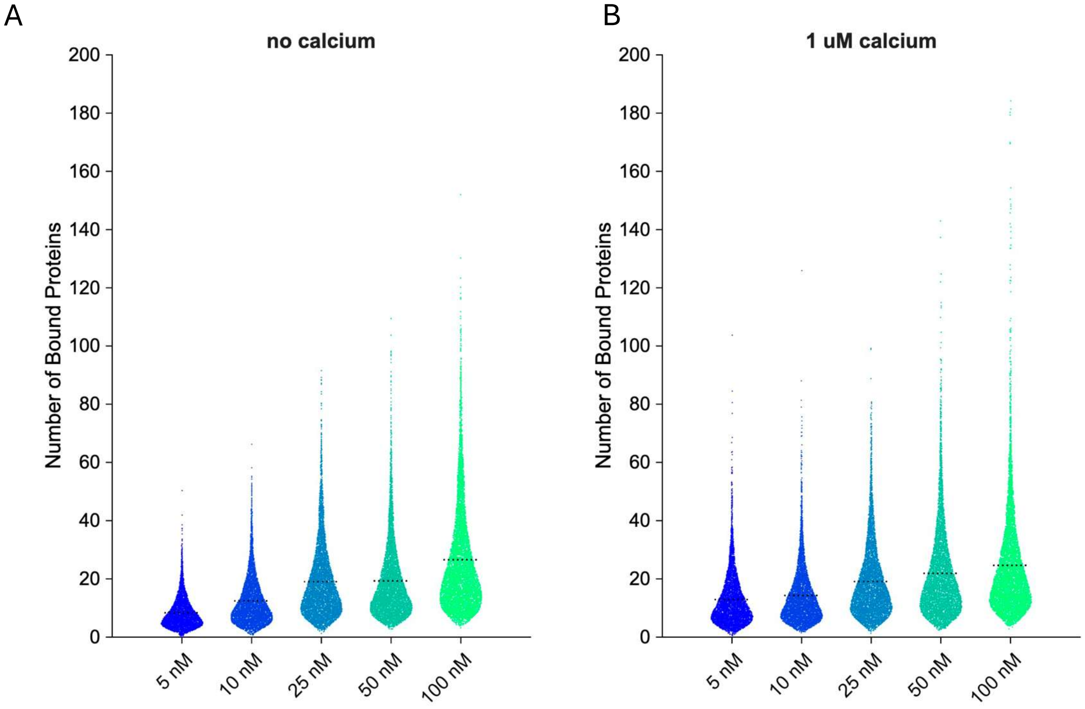


**Figure S9 – Number of bound clathrin triskelia per vesicle**

The raw data for the binding curves depicted in Figure 3B. Vesicles were composed of 88.5% DOPC, 10% DGS-NTA-$NI^{2+}$, 1% DPEG-15-biotin, and 0.5% ATTO-647N, and extruded through 200 nm pores. Each dot represents a vesicle for increasing concentrations of clathrin for **(A)** buffer conditions without $Ca^{2+}$ **(B)** 1 µM $Ca^{2+}$. The dashed line represents the mean.

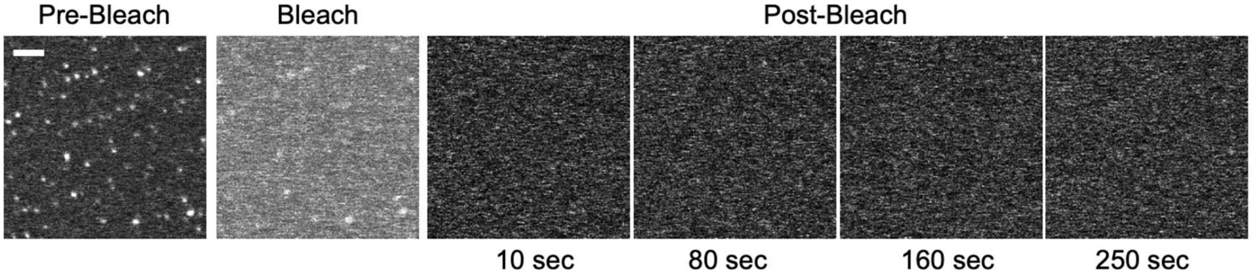


**Figure S10 – Bound clathrin triskelia do not undergo dynamic exchange on vesicles**

Representative fluorescence recovery after photobleaching (FRAP) image sequence of his-clathrin bound to vesicles. Vesicles were composed of 88.5 mol% DOPC, 10 mol% DGS-NTA-$NI^{2+}$, 1 mol% DPEG-15-biotin, and 0.5 mol% ATTO-647N, and synthesized via extrusion through 200 nm pores. Scale bar = 2 μm.

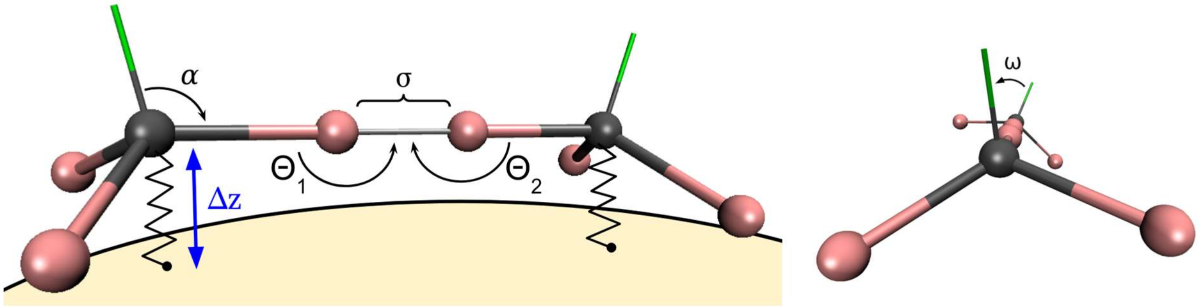


**Figure S11 – Coarse Grained clathrin model**

Left: Schematic diagram of two bonded coarse-grained (CG) rigid-body clathrin triskelia. Clathrin-binding sites are shown in pink. The triskelion body-fixed normal vector is shown as green. The indicated quantities are triskelion pucker angle α, bond length σ, bond angles θ, and triskelion-membrane bond length as Δz. Right: The final bond parameter is the normal vector dihedral angle ω. Equilibrium values are σ = 5 nm, θ = π rad, ω = 0, and Δz = 15 nm. The Δz force constant is fixed at 1000 $kT/nm^2$, while the others are varied between 200 and 4000 ($kT/nm^2$ or $kT/rad^2$) in order to tune lattice rigidity.

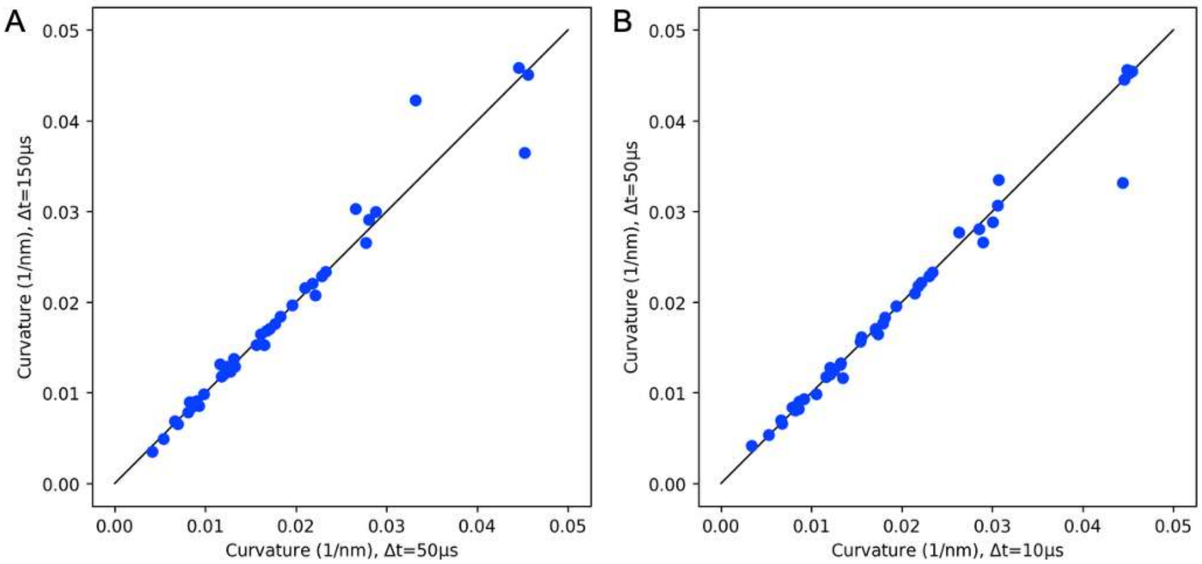


**Figure S12 – In Brownian Dynamics simulations, the degree of curvature generation does not change as we increase the rate that clathrin triskelia are added to the growing lattice.**

**(A)** The x-axis has a faster rate of clathrin addition to the lattice, with one per $\Delta t$ = 50 μs, vs a slower addition of once per $\Delta t$ = 150 μs on the y-axis. **(B)** The faster rate of addition with $\Delta t$ = 10 μs still produces a curvature consistent with slow lattice growth, demonstrating that the lattice growth rate is not limiting the degree of membrane invagination. The identity line is plotted for reference. Data are from the same parameter values used in Fig. 4b-d of the main text.

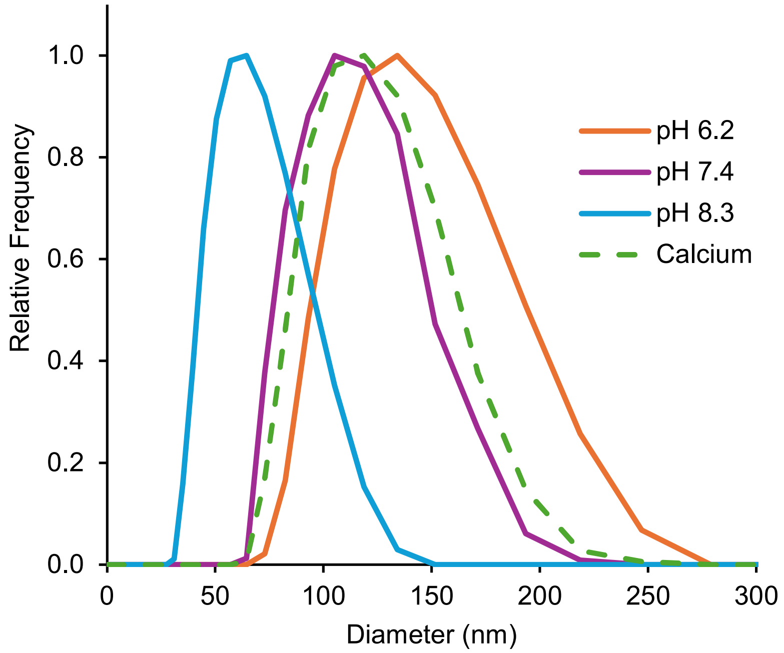


**Figure S13 – Clathrin Cage Diameters measured by DLS**

Hydrodynamic size distributions of clathrin from dynamic light scattering at pH 6.2, 7.4, 8.3, and with 1 μM $Ca^{2+}$ (pH 7.4). Data was used to generate Figure 4E.

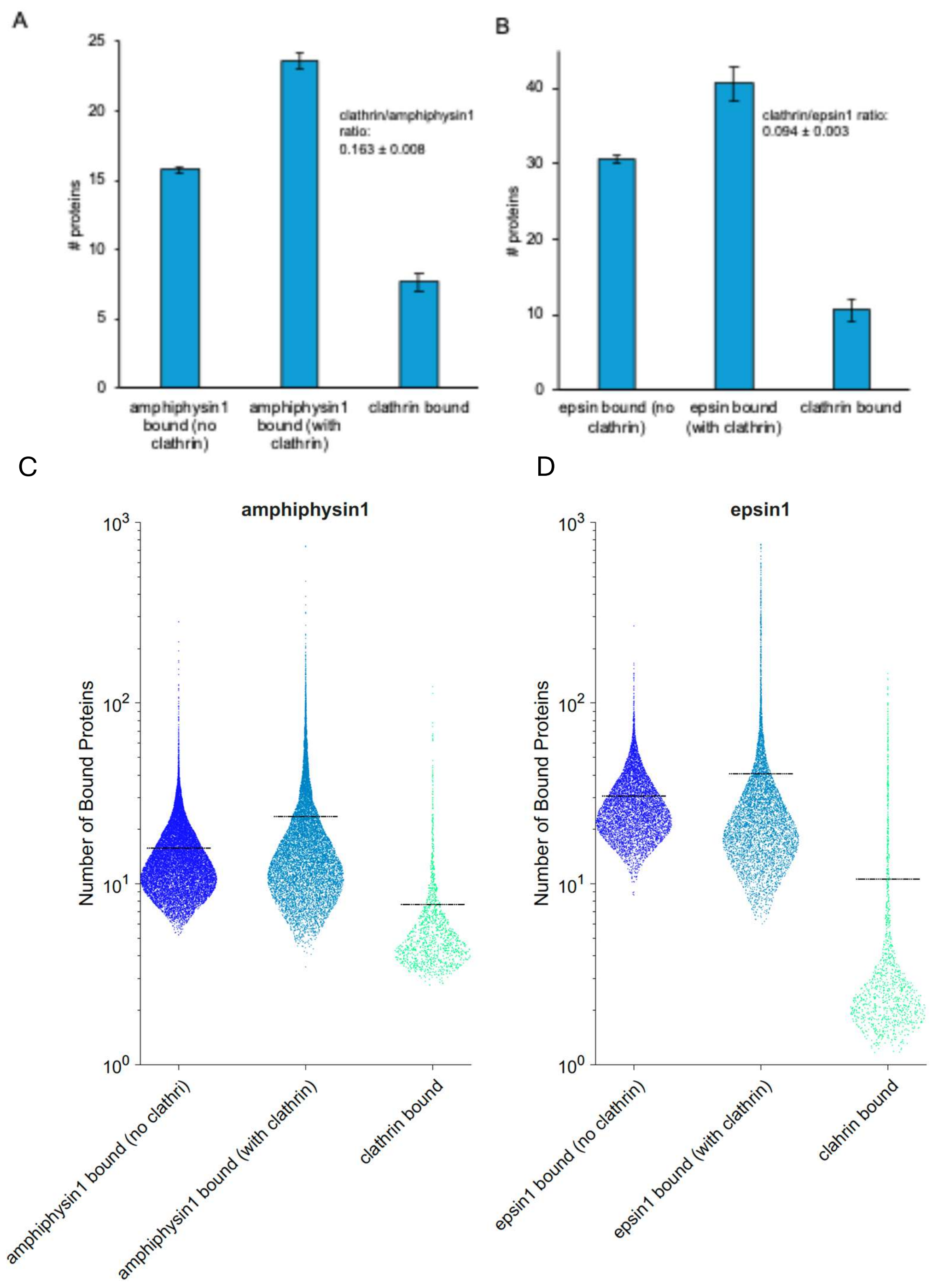


**Figure S14 – Number of bound adaptor proteins and clathrin per vesicle**

**(A)** Average number of amphiphysin1 (labeled Alexa-594) proteins bound to vesicles (composition: 78.5% DOPC, 5% PIP2, 15% DOPS, 1% DPEG-15-biotin, 0.5% DPPE-ATTO-488) in the presence and absence of clathrin (labeled Alexa-647). The average clathrin:amphiphysin1 ratio (i.e. number of clathrin triskelia bound per amphiphysin1) with 99% confidence interval is inlaid. Error bars represent 99% confidence interval. Amphiphysin1 concentration was 500 nM, clathrin concentration was 50 nM. **(B)** Average number of epsin1 (labeled Alexa-594) proteins bound to vesicles (composition: 73.5% DOPC, 10% PIP2, 15% DOPS, 1% DPEG-15-biotin, 0.5% DPPE-ATTO-488) in the presence and absence of clathrin (labeled Alexa-647). The average clathrin:epsin1 ratio (i.e. number of clathrin triskelia bound per epsin1) with 99% confidence interval is inlaid. Error bars represent 99% confidence interval. Epsin1 concentration was 2 μM, clathrin concentration was 50 nM. **(C)** Raw data used to generate **(A)** and **(D)** is the raw data used to generate **(B)**. Each dot represents a vesicle, and the dashed line represents the mean.
